# Branching of sporogenic aerial hyphae in *sflA* and *sflB* mutants of *Streptomyces coelicolor* correlates to ectopic localization of DivIVA and FtsZ in time and space

**DOI:** 10.1101/2020.12.26.424426

**Authors:** Le Zhang, Joost Willemse, Paula Yagüe, Ellen de Waal, Dennis Claessen, Gilles P. van Wezel

## Abstract

Bacterial cytokinesis starts with the polymerization of the tubulin-like FtsZ, which forms the cell division scaffold. SepF aligns FtsZ polymers and also acts as a membrane anchor for the Z-ring. While in most bacteria cell division takes place at midcell, during sporulation of *Streptomyces* many septa are laid down almost simultaneously in multinucleoid aerial hyphae. The genomes of streptomycetes encode two additional SepF paralogs, SflA and SflB, which can interact with SepF. Here we show that the sporogenic aerial hyphae of *sflA* and *sflB* mutants of *Streptomyces coelicolor* frequently branch, a phenomenon never seen in the wild-type strain. The branching coincided with ectopic localization of DivIVA along the lateral wall of sporulating aerial hyphae. Constitutive expression of SflA and SflB largely inhibited hyphal growth, further correlating SflAB activity to that of DivIVA. SflAB localized in foci prior to and after the time of sporulation-specific cell division, while SepF co-localized with active septum synthesis. Foci of FtsZ and DivIVA frequently persisted between adjacent spores in spore chains of *sflA* and *sflB* mutants, at sites occupied by SflAB in wild-type cells. This may be caused by the persistance of SepF multimers in the absence of SflAB. Taken together, our data show that SflA and SflB play an important role in the control of growth and cell division during *Streptomyces* development.

## INTRODUCTION

Streptomycetes are multicellular mycelial bacteria that reproduce via sporulation (Claessen *et al*., 2014; Flärdh and Buttner, 2009). As producers of half of all known antibiotics as well as many anticancer, antifungal and immunosuppressant compounds, streptomycetes are of great medical and biotechnological importance (Barka *et al*., 2016; Hopwood, 2007). The mycelial life style of streptomycetes imposes specific requirements for the control of growth and cell division (Jakimowicz and van Wezel, 2012; McCormick, 2009), and they have an unusually complex cytoskeleton (Bagchi *et al*., 2008; Celler *et al*., 2013).

The *dcw* gene cluster contains various genes required for division and cell wall synthesis (Tamames *et al*., 2001; Vicente and Errington, 1996). Some genes in this cluster have gained species-specific functions. An obvious example is DivIVA, which in *Bacillus subtilis* is involved in division-site localization by preventing accumulation of the cell division scaffold protein FtsZ (Marston *et al*., 1998b), while in Actinobacteria DivIVA is required for apical growth (Flärdh, 2003). As a consequence, *divIVA* is dispensable in *B. subtilis* but essential for growth in Actinobacteria (Flärdh, 2003; Letek *et al*., 2008). Conversely, *ftsZ* is essential in *B. subtilis*, but is no required for normal growth of Actinobacteria (*McCormick et al., 1994*).

The control of cell division is radically different between the mycelial streptomycetes and the planktonic *Bacillus subtilis*, which is perhaps not surprising due to the absence of a defined mid-cell position in the long hyphae of streptomycetes. In rod-shaped bacteria, many proteins have been identified that assist in septum-site localization, such as FtsA and ZipA (Hale and de Boer, 1997; Pichoff and Lutkenhaus, 2002; RayChaudhuri, 1999) and ZapA (Gueiros-Filho and Losick, 2002). Septum-site localization is negatively controlled, via the action of Min, which prevents Z-ring assembly away from mid-cell (Marston *et al*., 1998a; Raskin and de Boer, 1997), and by nucleoid occlusion that prevents formation of the Z-ring over non-segregated chromosomes (Bernhardt and de Boer, 2005; Woldringh *et al*., 1991; Wu and Errington, 2004, 2012). Direct homologs of any of these control proteins are missing in streptomycetes.

Streptomycetes have two different mechanisms of cell division. During vegetative growth, divisome-independent cell division occurs, whereby occasional cross-walls separate the vegetative hyphae into connected multicellular compartments. The cross-walls depend on FtsZ, but not on other canonical divisome proteins such as FtsI, FtsL and FtsW (Jakimowicz and van Wezel, 2012; McCormick, 2009; Mistry *et al*., 2008). Interestingly, mutants lacking *ftsZ* are viable, forming long hyphae devoid of septa (McCormick et al., 1994). Intricate membrane assemblies ensure that chromosome-free zones are created during septum formation in vegetative hyphae, apparently protecting the DNA from damage during division (Celler *et al*., 2016; Yagüe *et al*., 2016). Reproductive and divisome-dependent cell division occurs exclusively in sporogenic aerial hyphae. Sporulation-specific cell division in *Streptomyces* may therefore be regarded as canonical cell division as it requires all components of the divisome. At the onset of sporulation, up to 100 septa are formed more or less simultaneously, see as spirals of FtsZ in the aerial hyphae. Cell division is positively controlled, via the direct recruitment of FtsZ by the membrane-associated SsgB (Willemse et al., 2011). SsgB is a member of the SsgA-like proteins, which only occur in morphologically complex actinomycetes (Jakimowicz and van Wezel, 2012; Traag and van Wezel, 2008). The localization of SsgB depends on the orthologous SsgA protein, which activates sporulation-specific cell division (Kawamoto *et al*., 1997; van Wezel *et al*., 2000).

Four genes lie between *ftsZ* and *divIVA* in the *dcw* cluster of streptomycetes, in the order *ftsZ-ylmD-ylmE-sepF-sepG-divIVA*. The small transmembrane protein SepG acts as an anchor for SsgB to the membrane and also controls nucleoid organization (Zhang et al., 2016). YlmDE form a likely toxin-antitoxin system, whereby YlmD acts as a toxin that has detrimental effects on sporulation-specific cell division (Zhang *et al*., 2018). SepF is involved in early division control by stimulating the polymerization of FtsZ. In *B. subtilis*, SepF forms large rings of around 50 nm in diameter in vitro, and assists in bundling of FtsZ filaments (Hamoen *et al*., 2006; Ishikawa *et al*., 2006). SepF interacts with the membrane via its N-terminal domain (Duman et al., 2013), and plays a role in both Z-ring assembly and anchoring. In the actinomycete *Mycobacterium* SepF also interacts with FtsZ, and is essential for division (Gola *et al*., 2015; Gupta *et al*., 2015). Thus, SepF is a rare example of a cell division control protein that is shared between firmicutes and by actinobacteria.

In this work, we analyzed the role of two paralogs of SepF in development and sporulation-specific cell division *Streptomyces coelicolor*. These are encoded by SCO1749 and SCO5967, which we designated *sflA* and *sflB* (for *sepF*-like), respectively. SflA and SflB play an important role in the control of development of the aerial hyphae, whereby branching spore chains were frequently seen in *sflA* and *sflB* mutants, coinciding with the unusual localization of DivIVA along the lateral wall and between spores. Conversely, overexpression of *sflA* or *sflB* resulted in reduced growth of the vegetative hyphae. FtsZ foci also persisted during spore maturation in *sflA* and *sflB* mutants. These data suggest that SflAB help to prevent the ectopic assembly of DivIVA and FtsZ during sporulation of *Streptomyces*.

## RESULTS

### Three *sepF*-like genes in *Streptomyces*

Three genes with homology to *sepF* were found on the *S. coelicolor* genome. The canonical *sepF* gene (SCO2079) lies within the *dcw* cluster in close proximity to *ftsZ*, an arrangement that is conserved in all Gram-positive bacteria. Two *sepF*-like (*sfl*) genes, *sflA* (SCO1749) and *sflB* (SCO5967), are located elsewhere on the *S. coelicolor* chromosome. SepF is a predicted 213 aa protein, while SflA (146 aa) and SflB (136 aa) are significantly smaller. Thus, SflA and SflB have lengths very similar to that of SepF of *Bacillus subtilis* (139 aa; accession number KFK80720). Alignment of the three proteins and their comparison to SepF of *B. subtilis* and *Mycobacterium smegmatis* is presented in Fig. 1; predicted α-helices and β-strands are boxed with dotted and solid lines, respectively. Compared to SflA and SflB, SepF proteins of *S. coelicolor* and *M. smegmatis* have an approximately 60 aa internal extension at the N-terminal half. The presence of three *sepF*-like genes is common in Actinobacteria, except for *Coriobacteriaceae*, which only have *sepF*. The N-terminal α-helix (aa 1-12) of *Bacillus* SepF is essential for lipid binding to support cell division (Duman et al., 2013). Based on the predicted secondary structure of the protein (using JPRED), this α-helix is absent in SflB (Cole et al., 2008), suggesting that this protein may not bind to the membrane. Conversely, the C-terminal domain of SepF, which is involved in the interaction with FtsZ (Duman *et al*., 2013; Gola *et al*., 2015; Gundogdu *et al*., 2011; Gupta *et al*., 2015), is conserved in SflA and SflB.

**Figure 1.**
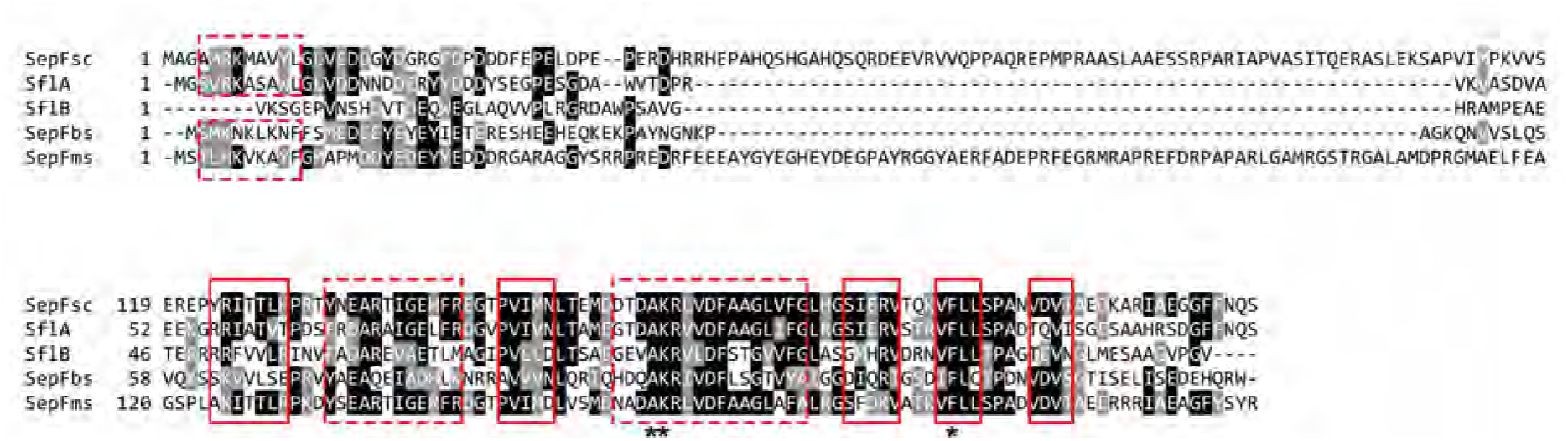
Alignment of SepF proteins. Amino acid sequences of SepF proteins from *B. subtilis* (SepFbs), *M. smegmatis* (SepFms) and *S. coelicolor* (SepFsc), and two SepF paralogs of *S. coelicolor* (SflA and SflB) were aligned using Boxshade program. Identical residues are shaded in black; conservative changes are shaded in grey. α-helices and β-strands in the predicted secondary structures (via JPRED) of are boxed by red dotted line and solid line, respectively. Essential amino acids for FtsZ interaction were highlighted with star.

### Deletion of *sflA* and *sflB* affects colony morphology

To analyze the role of SflA and SflB in growth and development of *Streptomyces*, deletion mutants were created for the two genes, separately and in combination, using a strategy based on the instable multi-copy plasmid pWHM3 (Swiatek et al., 2012). Briefly, the +10/+426 section of *sflA* or the +10/+356 region of *sflB* (relative to the start of the respective genes) was replaced by the apramycin resistance cassette, which was subsequently removed using the Cre-*lox* system, leaving only the scar sequence, thereby generating an in-frame deletion mutant (see Materials and Methods). The *sfl* single and double mutants sporulated well on SFM agar plates, developing abundant grey-pigmented spores after 3 days of growth, suggesting that these proteins are dispensable for sporulation (Fig. 2B). Nonetheless, the timing of development was mildly affected in the mutants. Deletion of *sflA* accelerated aerial growth and sporulation, while deletion of *sflB* delayed sporulation. In *sflAB* double mutants, aerial hyphae formation was accelerated while sporulation was delayed (Fig. S1).

**Figure 2.**
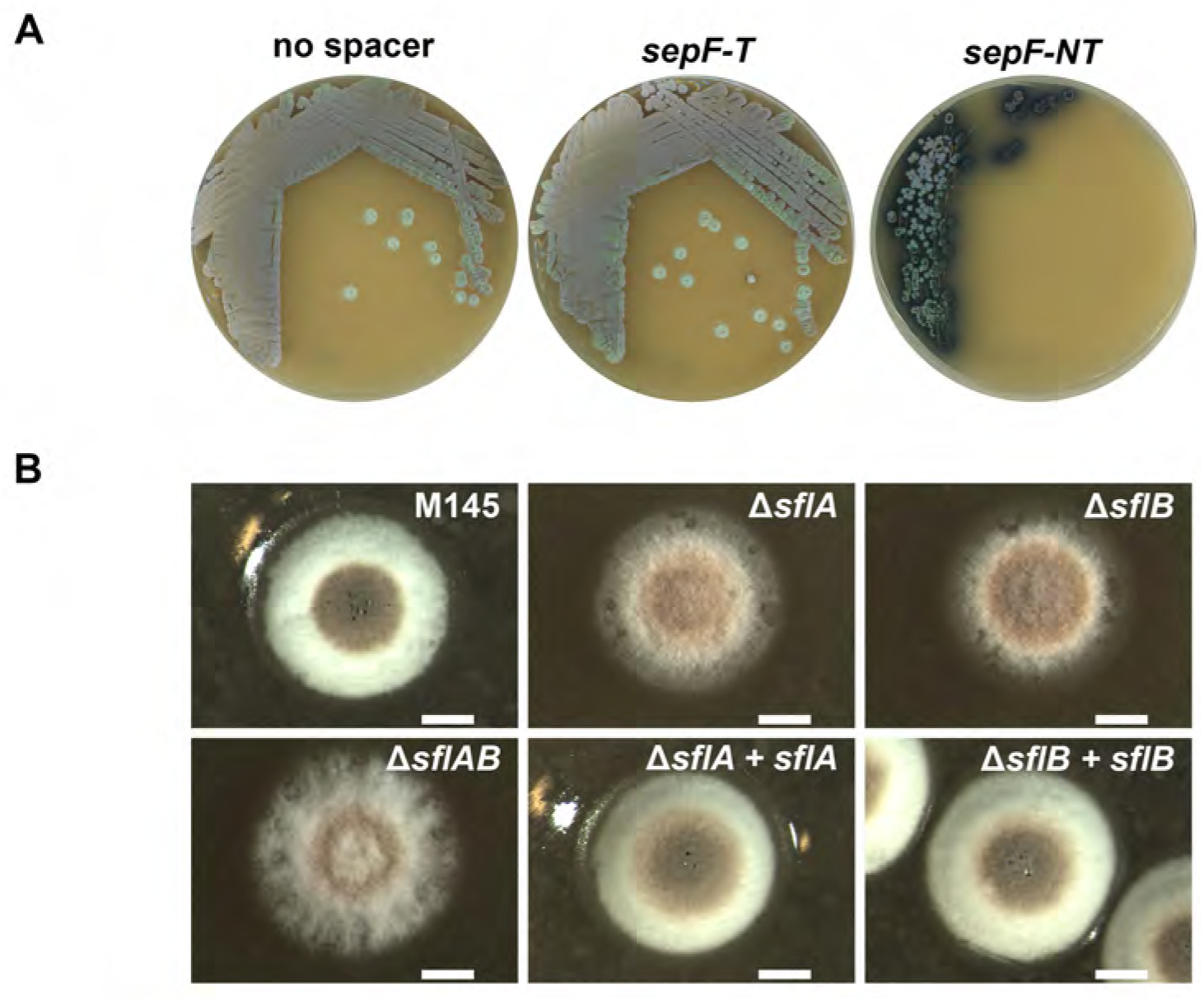
Phenotypic analysis of *sepF* and *sfl* mutants. *sepF* knockdown mutant shows severe developmental defect when the spacer in CRISPRi system targets non-template strand **(A)**. Stereomicrographs show representative colonies of *S. coelicolor* M145, its *sflA* and *sflB* null mutants and complemented strains. Strains were grown on SFM agar plates for three days at 30°C. Note that colonies of *sfl* mutants were ‘fluffier’ than those of the parental strain M145, and expression of wild-type SflA or SflB restore smooth colony edge to the corresponding mutants Bar, 1 mm **(B)**.

Interestingly, while *S. coelicolor* M145 formed colonies with a smooth edge, those of *sflA* or *sflB* mutants had a ‘fluffier’ phenotype, a difference that was more pronounced in *sflAB* double mutants (Fig. 2B). Genetic complementation of *sflA* and *sflB* null mutants by the introduction of plasmids pGWS1005 (expressing *sflA* from the *ftsZ* promoter) and pGWS1006 (expressing *sflB* from the *ftsZ* promoter), respectively, restored the wild-type colony morphology. This indicates that the abnormal colony morphology of the mutants was indeed due to the deletion of the *sfl* genes. To investigate the change in colony morphology in *sfl* mutants, the tip-to-branch distance was measured in young vegetative hyphae that had been grown for 20 h. This average tip-to-branch distance was 15.05 ± 5.14 μm in the parental strain M145, while it had increased significantly in *sflA, sflB* and *sflAB* mutants, where the distance was 19.79 ± 9.15 μm, 18.84 ± 9.06 μm and 19.89 ± 7.12 μm, respectively (*p* < 0.001). The longer tip-to-branch distance in *sfl* mutants - and thus reduced compactness of the mycelia - may explain the altered colony morphology of *sfl* mutants.

We also attempted to delete *sepF*, but failed to do so despite many attempts. Therefore, CRISPRi was employed to knockdown *sepF* and obtain insights into its possible functional linkage to *sflAB*. The CRISPRi system we used was modified from pCRISPR-dCas9 (Tong et al., 2015) by expressing Cas9 from the constitutive *gapdh* promoter, using vector pSET152 that integrates at the øC31 attachment site on the *S. coelicolor* chromosome (see Materials and Methods section for details)(Ultee et al., 2020). Introduction of control constructs pGWS1050 and pGWS1353, which contain either no spacer or a spacer targeting the template strand of *sepF*, respectively, did not affect growth or development of *S. coelicolor* (Fig. 2A). Conversely, introduction of pGWS1354, which carries a spacer targeting the non-template strand of *sepF*, into *S. coelicolor* M145, resulted in severe developmental defects and overproduction of actinorhodin (Fig. 2A). Transmission electron microscopy (TEM) showed that vegetative hyphae wherein *sepF* was knocked down using CRISPRi lacked cross-walls (Fig. S2). The phenotype of *sepF* mutants was very similar to that reported for *ftsZ* null mutants (McCormick et al., 1994), in line with the expected crucial role of SepF in Z-ring formation in *S. coelicolor*. The severe phenotype of the *sepF* knock-down mutants suggests that *sflA* and *slfB* cannot functionally compensate for the lack of *sepF*.

### Sporogenic aerial hyphae of *sflA* and *sflB* null mutants show unusual branching

Surface-grown *S. coelicolor* M145 and its *sflA, sflB* and *sflAB* mutants were analyzed in more detail by cryo-scanning electron microscopy (SEM). After three days of growth, *S. coelicolor* M145 produced abundant and regular spore chains (Fig. 3A). However, strains lacking *sflA (ΔsflA* and *ΔsflAB*) produced fewer spore chains (Fig. 3B & 3D), while deletion of only *sflB* did not significantly affect sporulation (Fig. 3C). Strikingly, sporogenic aerial hyphae of *sflA, sflB* and *sflAB* null mutants branched frequently (Fig. 3E-G), a phenotype that was never seen in the wild-type strain. Introduction of wild-type copies of *sflA* or *sflB* into the respective mutants largely complemented the mutant phenotypes, and prevented branching (Fig. 3H-I). Some variability in spore sizes was still observed, perhaps as the result of a difference in expression level of the proteins from the chromosomal and from the plasmid-borne genes.

**Figure 3.**
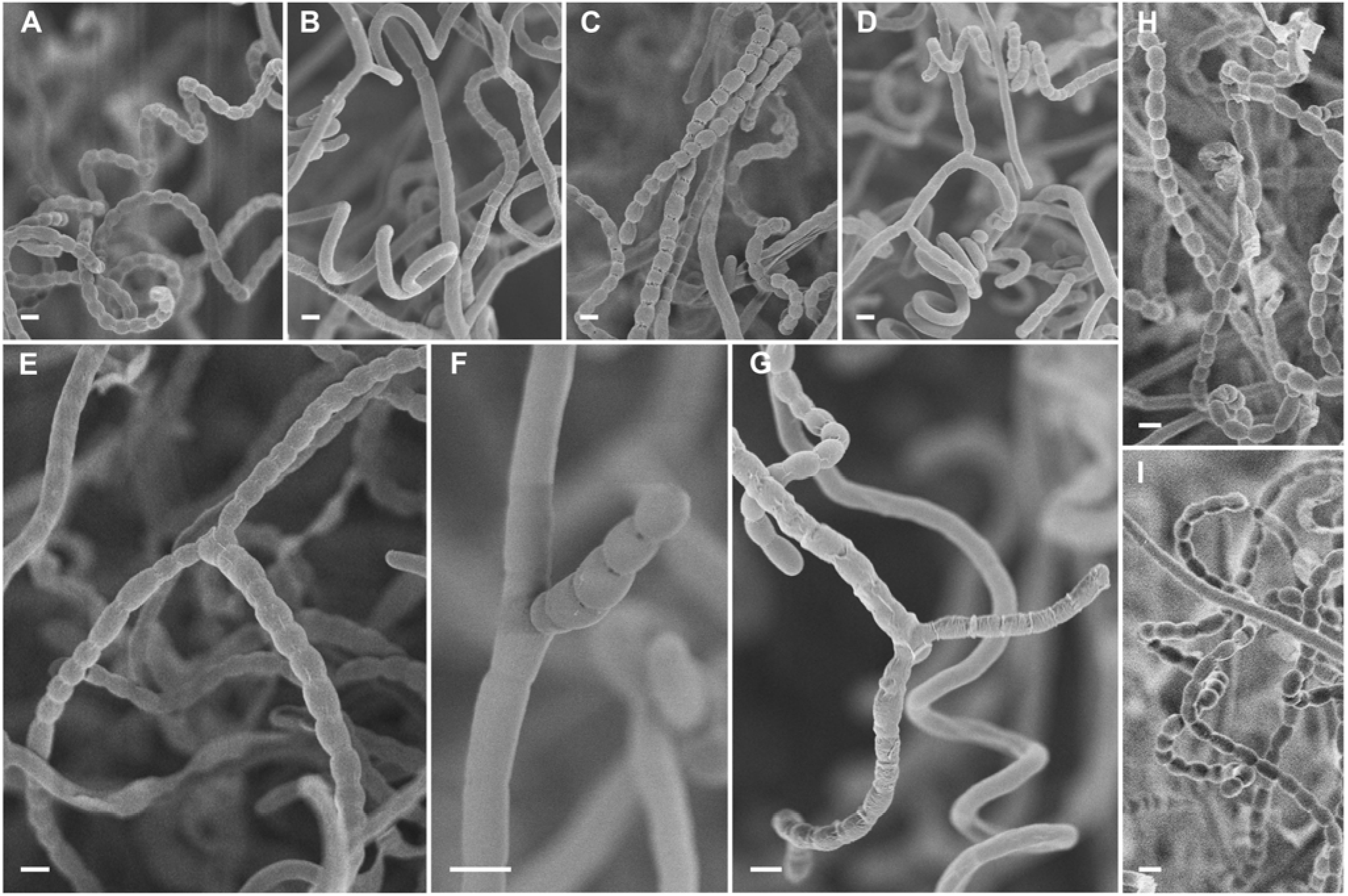
Cryo-scanning electron micrographs of spore chains of *S. coelicolor* M145, its *sfl* mutants and complemented *sfl* mutants. Wild-type *S. coelicolor* M145 **(A)** sporulated abundantly after three days of incubation, while mutants lacking either *sflA* **(B & E)** or *sflAB* **(D & G)** showed reduced sporulation; the *sflB* null mutant **(C & F)** produced comparable amount of spores as the parental strain. Most notable change in all mutants was that the spore chains frequently branched, while spore chains in genetically complemented *sflA* **(H)** and complemented *sflB* **(I)** did not show any branching. Cultures were grown on SFM agar plates for 5 days at 30°C. Bar, 1 μm.

Transmission electron microscopy (TEM) was used to image thin sections at high resolution. This again revealed branching spore chains in *sflA* and *sflB* mutants (Fig. 4, arrows) and variation in spore sizes. Furthermore, while wild-type spores had a typical dark (electron-dense) spore wall and well-condensed DNA, the spores of the mutants typically had lighter (electron-lucent) spore walls as well as less clearly visible DNA in many of the spores (Fig. 4 B-D). This suggests pleiotropic changes in spore morphogenesis and maturation in *sfl* genes mutants. As was already apparent from the SEM imaging, introduction of *sflA* and *sflB* into *sflA* and *sflB* null mutants, respectively, prevented branching of the spore chains, although the spore walls were still relatively thin (Fig. 4 E-F).

**Figure 4.**
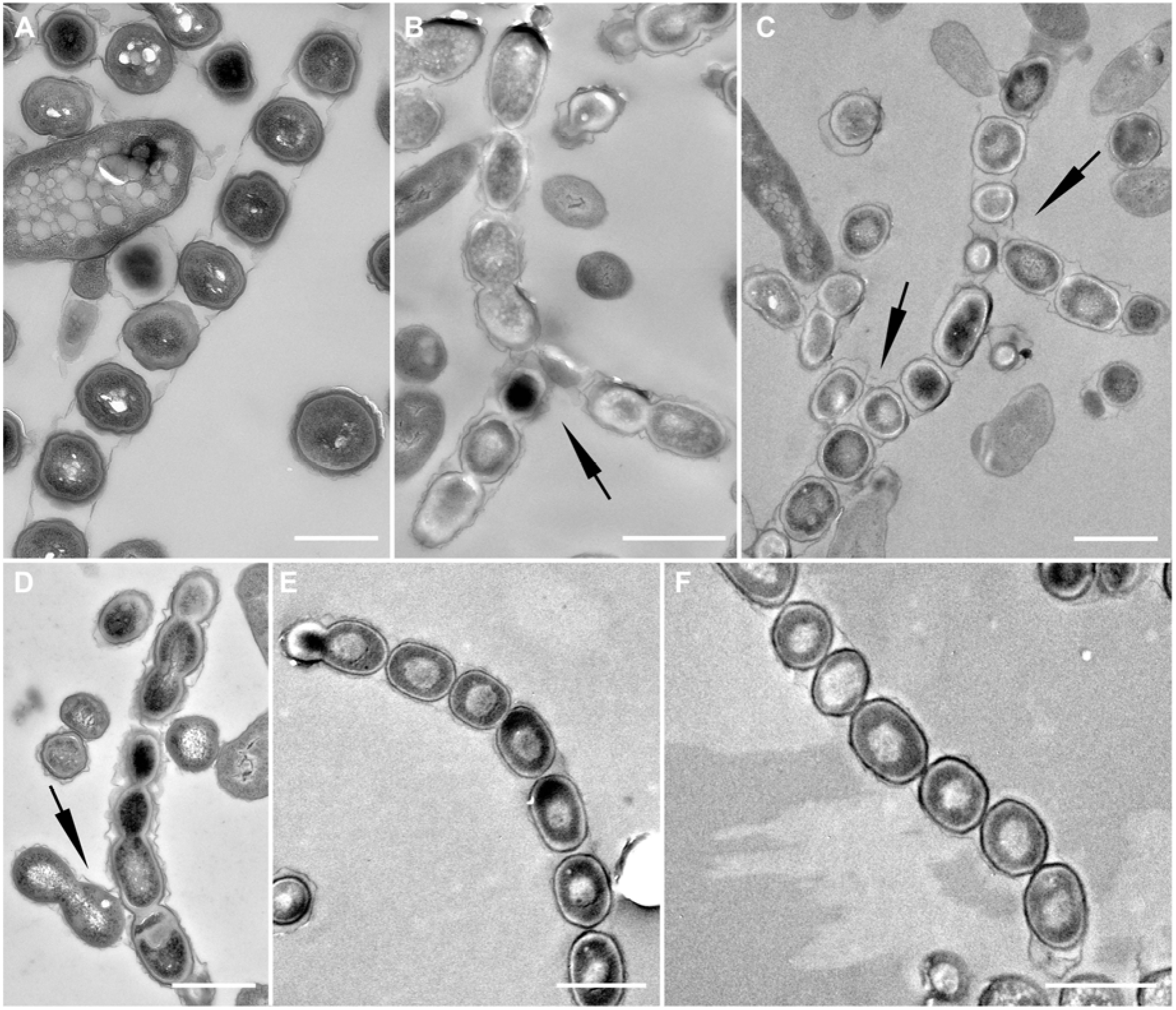
Transmission electron micrographs of spore chains of *S. coelicolor* M145, its *sfl* mutants and complemented *sfl* mutants. While spore chains of wild type M145 **(A)** do not branch and contain regularly sized spores, mutant lacking either *sflA* **(B)**, *sflB* **(C)** or *sflAB* **(D)** produce irregular spores and spore chains frequently branch, in line with the SEM images (Figure 3). Complemented *sflA* **(E)** and complemented *sflB* **(F)** produced unbranched spore chain as wild type. Cultures were grown on SFM agar plates for 5 days at 30°C. Arrows indicate branching points of spore chains. Bar, 1 μm.

### Effect of enhanced expression of the *sepF* and *sfl* genes

To study the effect of overexpression of SepF paralogs in *S. coelicolor*, the *sflA, sepF* and *sflB* genes were all cloned individually behind the *ermE* promoter region, which encompasses a strong constitutive promoter and an optimized ribosome binding site (see Materials and Methods for details), and the expression cassettes were then inserted in the multi-copy shuttle vector pWHM3. The expression constructs were designated pGWS774, pGWS775 and pGWS776, respectively. pWHM3 is an unstable plasmid that is easily lost and its copy number largely depends on the level of thiostrepton (van Wezel et al., 2005). The thiostrepton concentration controls the copy number of pWHM3, with copy number proportional to the thiostrepton concentration.

Plasmids pGWS774 (expressing *sflA*), pGWS775 (*sepF*), pGWS776 (*sflB*) or control plasmid pWHM3 without insert were introduced into *S. coelicolor* M145. The transformants were then plated onto SFM agar plates with different concentrations of thiostrepton and the colony morphology investigated after 7 days of incubation (Fig. 5). On SFM media, even in the absence of thiostrepton, colonies overexpressing SflA (GAL44) or SflB (GAL46) were smaller than those of transformants harboring the empty plasmid (GAL70) or transformants over-expressing SepF (GAL45) (Fig. 5). In the presence of thiostrepton (20 μg/mL), the size of colonies over-expressing SflA or SflB were reduced further. Interestingly, spores of SflA- and SflB-overexpressing strains could be easily removed from the plates with a toothpick, leaving “clean” plates, suggesting they had lost the ability to attach to and invade into the agar surface (Fig. 5, third row). Conversely, SepF-overexpressing colonies still grew into agar, and the mycelia remained firmly attached to the plates (Fig. 5, third row). When the thiostrepton concentration was increased further to 50 μg/mL, colonies of transformants with SflA or SflB expression constructs were very tiny and irregularly shaped, while those with control plasmid or harboring the SepF expression construct were barely affected (Fig. 5). On R5 agar plates, similar tiny colonies were observed for SflA and SflB-overexpressing strains, whereby the colonies more or less ‘floated’ on the agar surface, showing severe developmental defect (Fig. S3).

**Figure 5.**
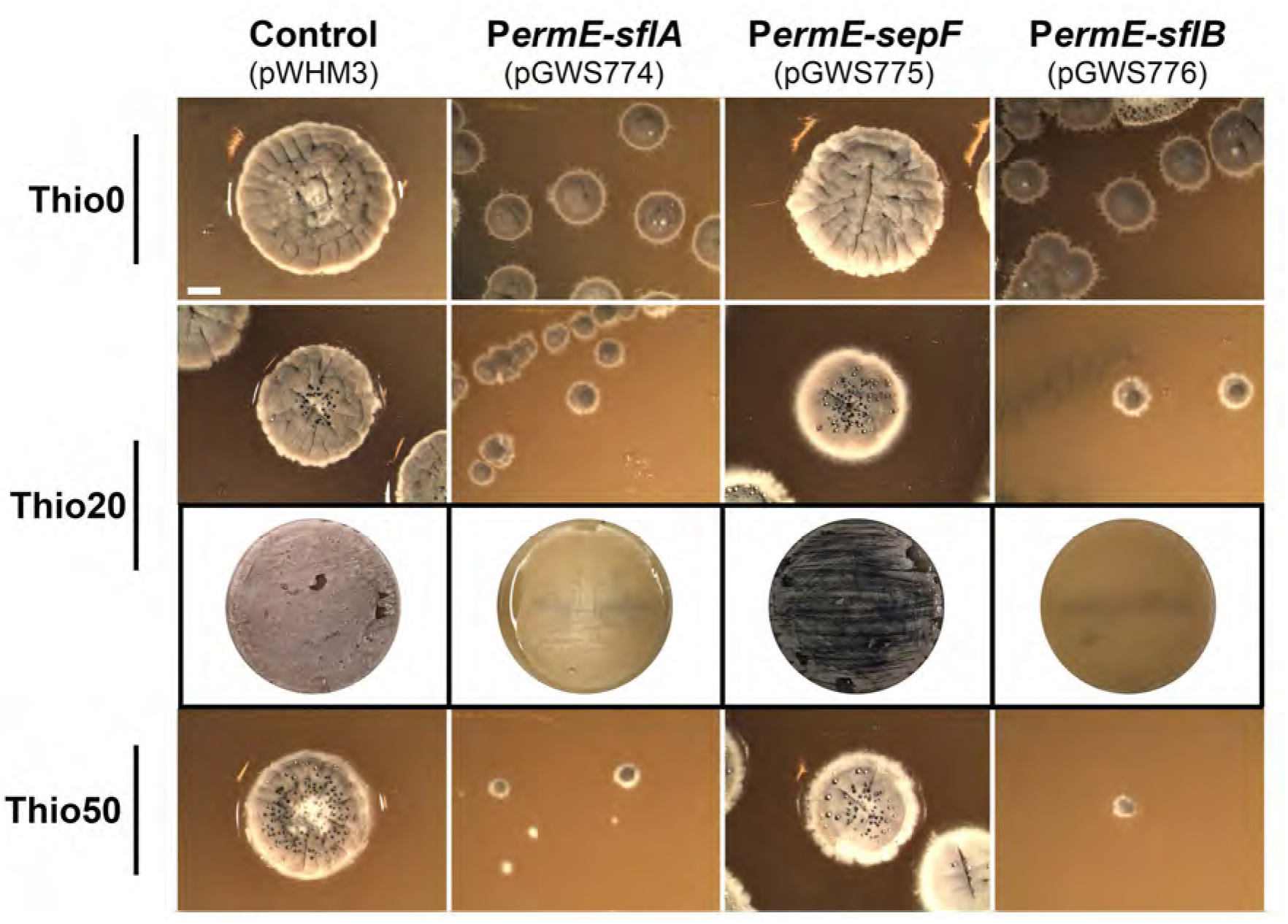
Effect of enhanced expression of *sepF* and *sfl* genes on colony morphology. Stereomicrographs showing the phenotype of GAL70 (*S. coelicolor* M145 + empty plasmid pWHM3 control), GAL44 (M145 + pGWS774, expressing *sflA*), GAL45 (M145 + pGWS775, expressing *sepF*) and GAL46 (M145 + pGWS776, expressing *sflB*) were grown on SFM plates containing different concentrations of thiostrepton (0-50 mg/ml). Plates were incubated for 7 days at 30°C. Over expression of *sflA* or *sflB* resulted in tiny colonies and no mycelium left on the plates after spore collection suggested the loss of attachment to agar, while overexpression of *sepF* didn’t affect colonial size and adherence. It should be noted that the tiny colonies produced by sflA or sflB overexpressing strains still show gray color, suggested that the sporulation were not inhibited. Bar, 2 mm.

To see if growth of the hyphae was affected, we analyzed young 9 h old vegetative hyphae. Interestingly, the hyphal length of control transformants carrying empty pWHM3 was 8.23 ± 3.57, while SflA- and SflB-overexpressing strains had a distance from germination site to hyphae tip of only 2.70 ± 1.59 μm and 2.70 ± 1.60 μm, respectively. The hyphal length of SepF-overexpressing colonies was less reduced, reaching on average 5.30 ± 1.70 μm. Taken together, we conclude that *sflA* or *sflB*, and to a lesser extent *sepF*, play a role in the control of tip growth.

### Altered localization of DivIVA and FtsZ in *sflA* and *sflB* mutants

Streptomycetes grow via extension of the hyphal tip, although the molecular mechanism of polar growth is still largely unknown (Jakimowicz and van Wezel, 2012). DivIVA is required for tip growth, whereby it localizes at apical sites and at new branches (Flärdh, 2003; Hempel *et al*., 2008). Therefore, DivIVA is a very good indicator for active tip growth, and we used this to study the onset of branching in the hyphae of wild-type and mutant strains. Construct pGWS800, harboring *Streptomyces venezuelae divIVA-egfp* under the control of its native promoter, was introduced into *sflA* and *sflB* null mutants. In wild-type cells, DivIVA-eGFP accumulated at tips of aerial hyphae, with 93% of the foci observed at apical sites. In aerial hyphae of *sflA* and *sflB* null mutants, DivIVA-eGFP foci were more widely distributed, not only at apical sites, but also along hyphae at the places without apparent branching, suggesting the emergence of new branching points (Fig. 6). In *sflA* and *sflB* mutants, 21% and 64% of the DivIVA-eGFP signals were observed along the lateral wall, respectively. Strikingly, DivIVA-eGFP localized abundantly in maturing spore chains of *sflA* and *sflB* mutants, while in wild-type spore chains no DivIVA-eGFP was observed (Fig. 6). The ectopic localization of DivIVA-eGFP in the absence of *sflA* or *sflB* suggests that their gene products play a role in the control of DivIVA localization and hence in determining apical growth of the hyphae in *Streptomyces*. This is consistent with the functional correlation of SflAB with tip growth and hyphal length.

**Figure 6.**
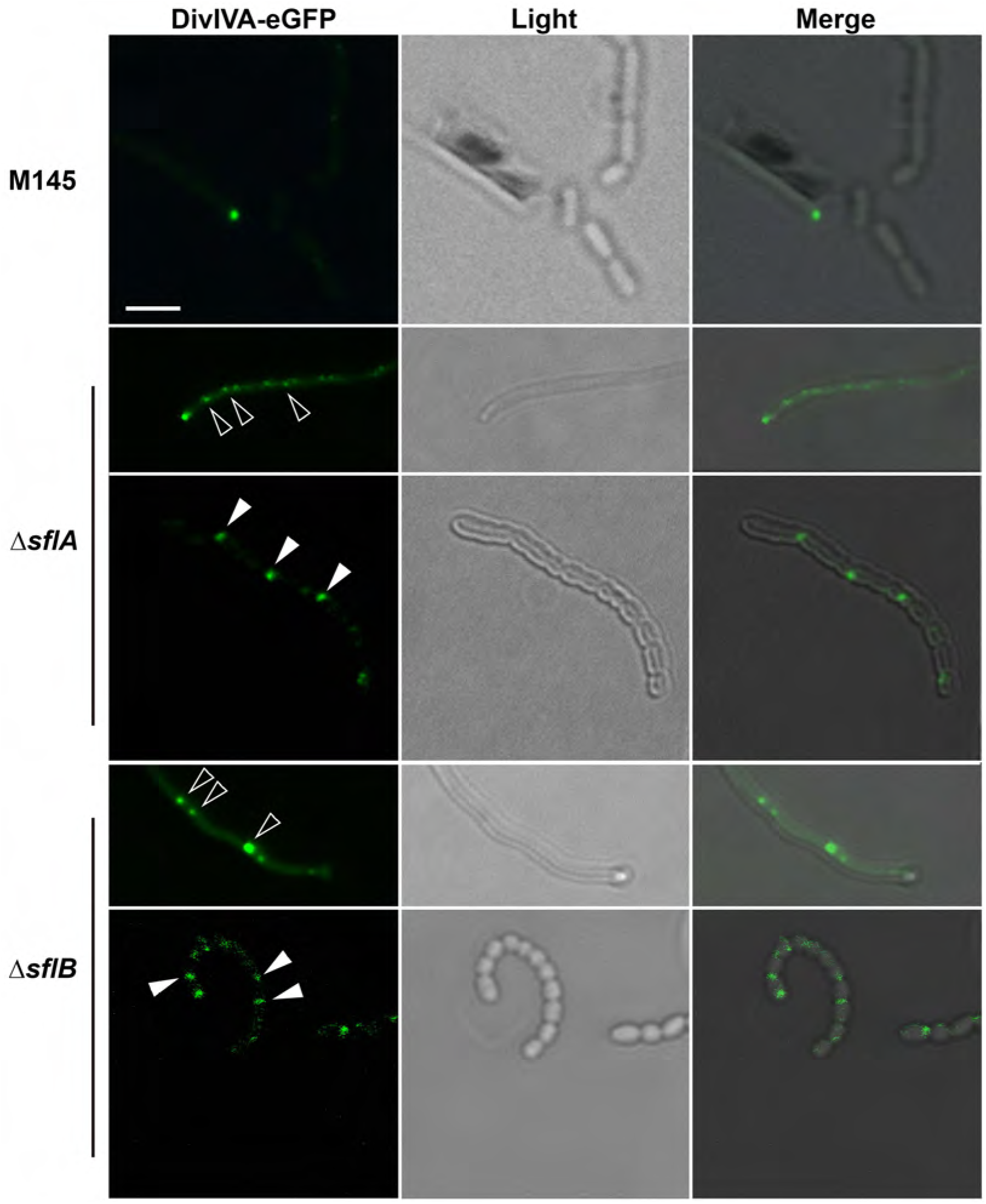
Localization of DivIVA-eGFP in *S. coelicolor* and its *sfl* null mutants. In aerial hyphae, DivIVA localized in wild-type cells mainly at tips while it was more dispersed in *sfl* mutants (indicated as empty arrow heads). DivIVA-eGFP was not detected in maturing spore chains of wild type cells, but it was often seen in that of *sfl* mutants (indicated as filled arrow heads). Bar, 2 μm.

To establish how FtsZ localizes in *sfl* mutants, construct pKF41 expressing FtsZ-eGFP (Grantcharova et al., 2005) was introduced into *S. coelicolor* and its *sflA, sflB* and *sflAB* mutants. In sporogenic aerial hyphae, FtsZ formed typical ladder-like patterns in all strains. Canonical Z-ladders were formed in *sfl* null mutants, although occasional misplaced septa were seen in *sflA* null mutants (Fig. 7, left). However, while FtsZ foci and rings disassembled and were absent in mature spore chains of wild-type *S. coelicolor*, they persisted in late sporogenic aerial hyphae of *sflA* and *sflB* mutants (Fig. 7, right). Prolonged Z-rings were observed in 46%, 28% and 72% of the premature spores of *sflA, sflB* and *sflAB* mutants, respectively, while they were not seen in wild-type spores. This corresponds very well to the ratios of incomplete septa in non-separated spores, which were 68% and 13% for *sflA* and *sflB* mutants, respectively, 79% for *sflAB* mutant and only 1% for the parent *S. coelicolor* M145. Taken together, the ectopic and continued localization of DivIVA and FtsZ in *sfl* null mutants throughout sporulation strongly suggests that SflA and SflB play an important role in controlling the dynamics of apical growth and cell division during *Streptomyces* development, and in particular ensure timely disassembly of DivIVA and FtsZ foci.

**Figure 7.**
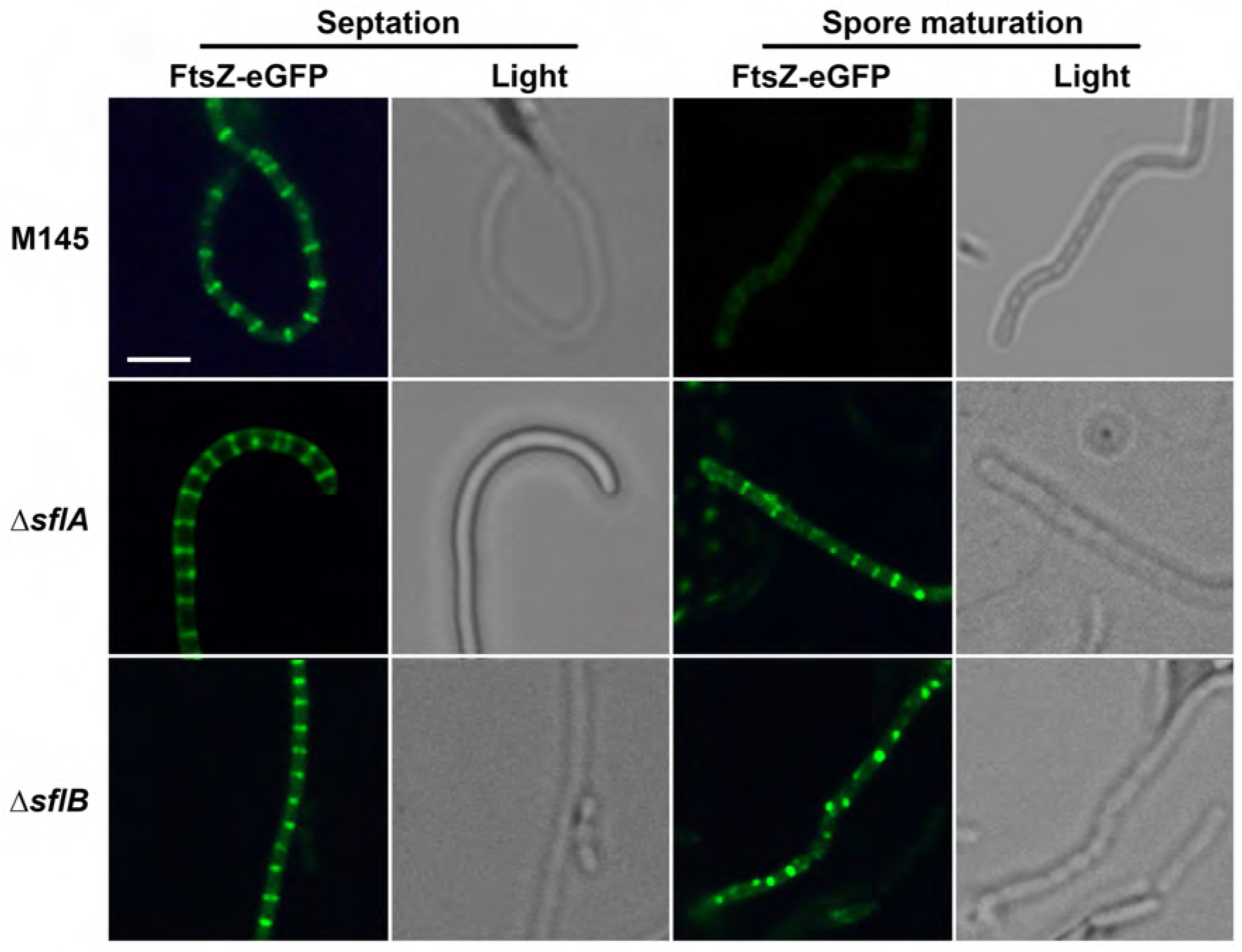
Localization of FtsZ-eGFP in *S. coelicolor* and its *sfl* null mutants. FtsZ forms ladder-like structure in sporogenic aerial hyphae of wild type and disappears in later developmental stage. While in *sfl* mutants FtsZ ladder remains longer even in spore maturing stage. Bar, 2 μm.

### Localization of SflA and SflB *in S. coelicolor*

To analyze the localization of the SepF paralogs, constructs were created in the integrative vector pSET152 containing either paralogue fused in frame behind *egfp* expressed from the *ftsZ* promoter region (see Materials and Methods). Constructs expressing eGFP-SflA, eGFP-SepF or eGFP-SflB from and were called pGWS784, pGWS785 and pGWS786 respectively. To analyze the colocalization of SepF and Sfl proteins, the gene for E2-Crimson was fused in frame with *sflA* and that for dTomato fused in frame with *sepF* (See Materials and Methods). The constructs expressing E2-Crimson-SflA or dTomato-SepF were named pGWS1380 and pGWS1383, respectively.

In young aerial hyphae, no specific localization of eGFP-SepF was observed prior to the onset of septum synthesis (Fig. 8A top row). Eventually, SepF-eGFP localized in a ladder-like pattern, similar to Z-ladders, which co-stained with the septa as seen by membrane staining using FM5-95 (Fig. 8A middle row). During spore maturation, no SepF-eGFP signal was detected (Fig. 8A bottom row). This indicates that SepF localizes in canonical fashion to sporulation septa, consistent with its role in Z-ring formation. Interestingly, eGFP-SflA and eGFP-SflB formed ring-like structures before septation had initiated (top row in Fig. 8B and 8C, respectively). When cell division had started, as visualized by membrane staining, eGFP-SflA and eGFP-SflB signals had largely disappeared (middle rows of Fig. 8B and Fig 8C, respectively). During spore maturation, when invagination between spores was clearly visible, the two proteins re-appeared at the junction between the adjacent spores (Fig. 8BC, bottom row). Thus, SflA and SflB localized specifically prior to and after the completion of septum synthesis, while SepF localized in rings primarily at the time when SflAB foci where no visible.

**Figure 8.**
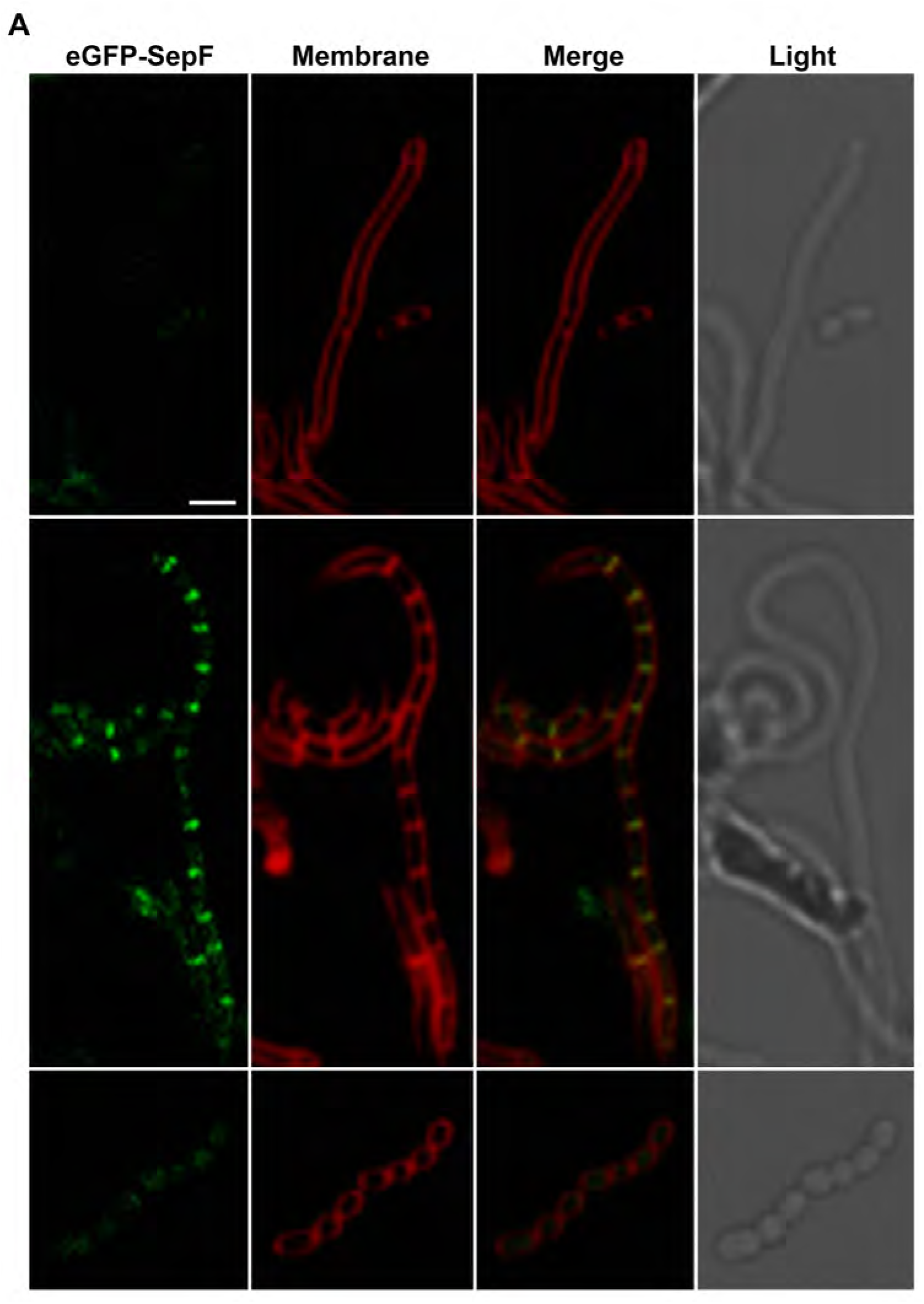
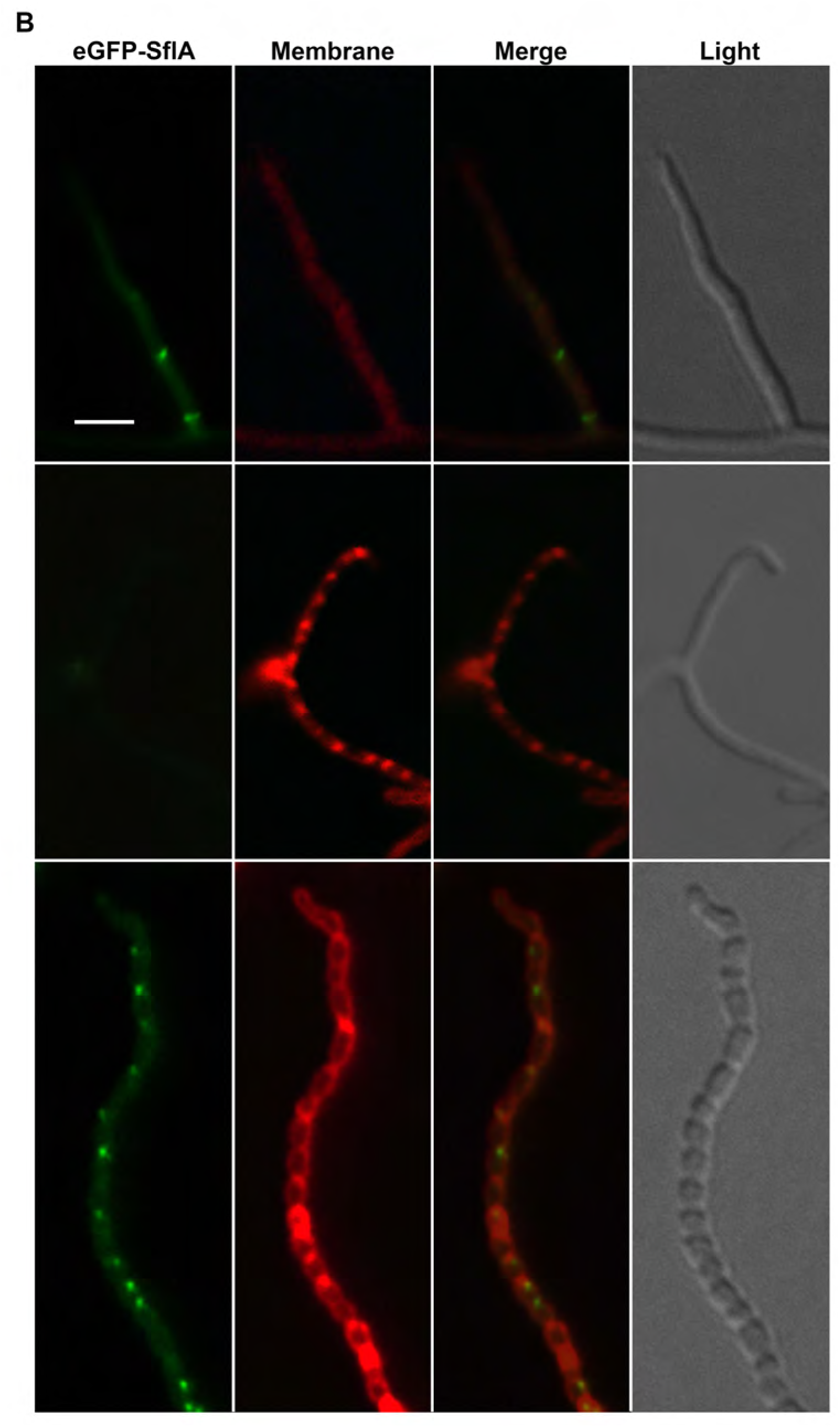
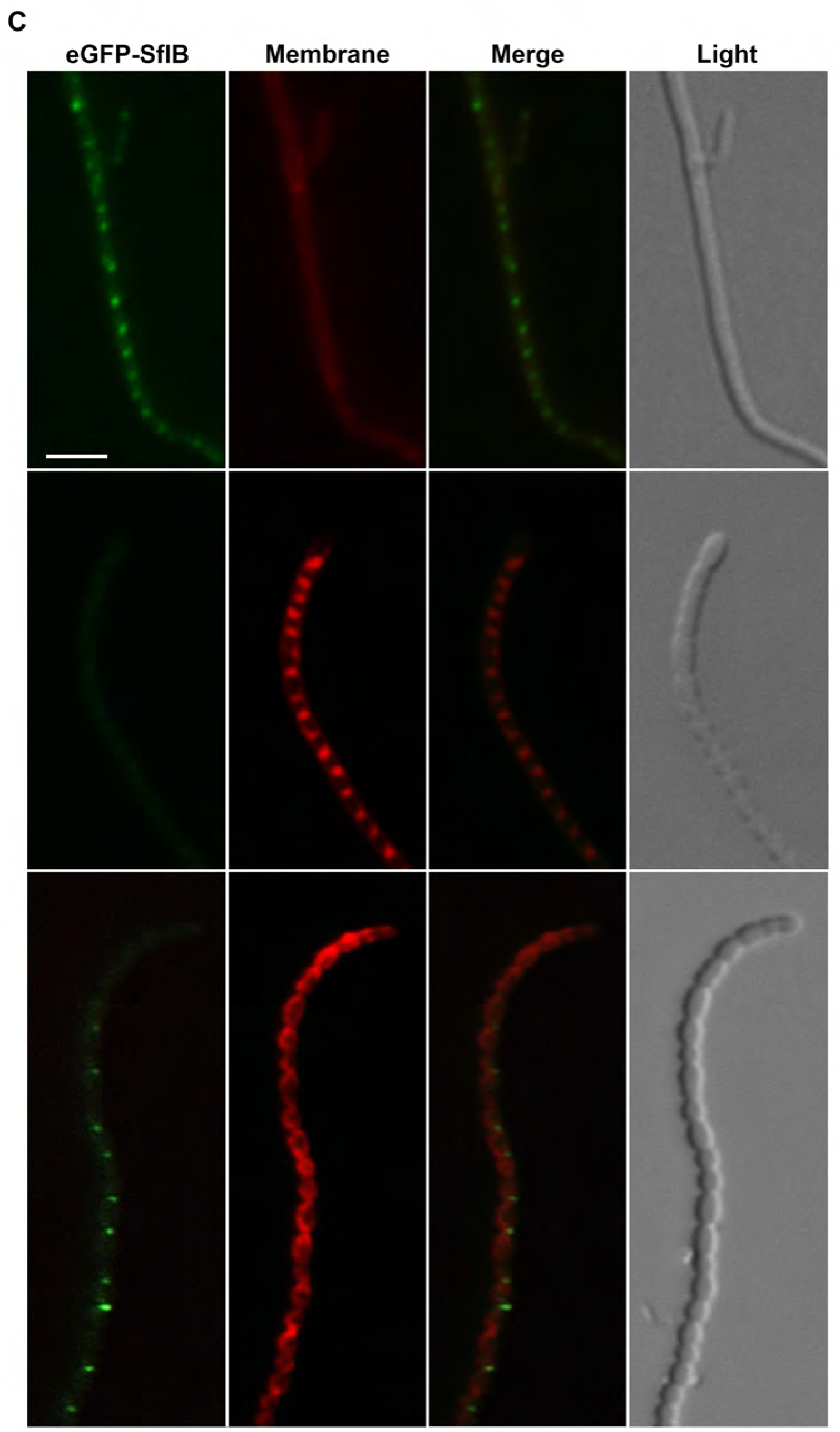
Localization of SepF, SflA and SflB in *S. coelicolor*. Fluorescence micrographs present three consecutive stages, namely prior to the onset of cell division (top panel), septum formation (middle panel) and spore maturation (bottom panel). Sporogenic aerial hyphae of *S. coelicolor* M145 were imaged by fluorescence microscopy visualizing the respective eGFP fusion proteins (green), membrane (stained with FM5-95; red) and corresponding light micrographs. As expected, eGFP-SepF (**A**) localizes in a ladder-like pattern that overlaps the sporulation septa. Foci of eGFP-SflA (**B**) forms foci along aerial hyphae prior to septum synthesis, re-appearing during spore maturation at invagination sites. Foci of eGFP-SflB (**C**) localize in a ladder-like pattern prior to septum synthesis, vanish as septal membranes formed and re-emerge during spore maturation. Bar, 2 μm.

The distinct localization patterns of SepF and Sfl proteins led us to investigate their colocalization. Indeed, SflA and SflB colocalized with each other, but most of the time they did not colocalize with SepF. However, on rare occasions, we did see colocalization between SepF and SflA or SflB, whereby they formed ring-like structures in sporogenic aerial hyphae (Fig. S5). This is consistent with experiments in *S. venezuelae*, which showed that both SepF and SflB (named SepF2 in *S. venezuelae*) colocalized with FtsZ, which indirectly confirmed the colocalization between SepF as SflB (Schlimpert et al., 2017). Live imaging of the sporulation process in solid-grown aerial hyphae is very difficult, due to the mobile nature of the airborne hyphae. We have been able to image the recruitment of FtsZ by SsgB, but this was during a short time frame and these are highly abundant proteins. Capturing the specific time when SflAB and SepF colocalize using live imaging was not feasible. Still, our results do show that while SepF and SflAB localized in differentially in terms of timing, there is a short time window when colocalization occurs. Dispersal of SflAB then marks the start of cell division.

## DISCUSSION

A major question in the developmental biology of *Streptomyces* that we seek to address is, how do *Streptomyces* ensure that septa are controlled in time and space in the long and multinucleoid hyphae? We have shown previously that in streptomycetes the correct localization of FtsZ is governed by a system of positive control, whereby the actinomycete-specific SsgA and SsgB proteins recruit FtsZ to the septum sites to initiate sporulation-specific cell division (Willemse et al., 2011). As a consequence, deletion of either *ssgA* or *ssgB* blocks sporulation (Keijser *et al*., 2003; van Wezel *et al*., 2000). Additionally, SepG (YlmG in *B. subtilis*) is an auxiliary protein that allows SsgB to dock to the membrane (Zhang et al., 2016). In this work we present a new piece of this jigsaw, which points at the possible existence of a layer of negative control during *Streptomyces* sporulation, revolving around the SepF-like proteins SflA and SflB.

The most eye-catching change in morphogenesis due to the deletion of either *sflA* or *sflB* was the extensive branching of the aerial hyphae and in particular of spore chains, which we have never seen in any of our wild-type streptomycetes. The tip-to-branch distance of vegetative hyphae was also extended in *sflA* and *sflB* null mutants, which likely contributes to the ‘fluffy’ morphology of the mutant colonies. Conversely, constitutive expression of SflA and SflB from the *ermE* promoter inhibited growth and reduced adhesion of the colonies to the agar surface, with as possible explanation that tip extension and hence also branching is impaired in the vegetative mycelium (Fig. 5 and Fig. S3). These data strongly suggest an inverse correlation between the expression level of SflA and SflB and polarisome activity. Indeed, we found mislocalization of DivIVA in *sflA* and *sflB* null mutants, with many foci along the lateral wall of aerial hyphae, instead of only apical localization. While in wild-type hyphae virtually all DivIVA-eGFP foci were located at the apex, in total 21% and 64% of the foci were observed along the lateral wall in *sflA* and *sflB* mutants, respectively. Since DivIVA drives tip growth and thus also branching, this likely explains the observed branching spore chains frequently observed in *sfl* null mutants (Fig. 3 and Fig. 4). The inhibition of growth following the constitutive expression of SflA or SflB in vegetative hyphae suggests that expression of these proteins throughout the life cycle directly or indirectly inhibits DivIVA during vegetative growth, which will then result in growth inhibition, as DivIVA is essential for tip growth.

Typical ladders of Z-rings were produced in young sporogenic aerial hyphae of both wild-type *S. coelicolor* and in *sflA* or *sflB* null mutants, though the distance between adjacent Z-rings in mutants varied more in the mutants. Importantly, besides for DivIVA, we also noticed strongly prologued and ectopic localization of FtsZ in mutants lacking *sflA* and/or *sflB*. While Z-ladders disappeared in mature spore chains of the parental strain *S. coelicolor* M145, ladders and foci persisted in the different *sfl* mutants during spore maturation, strongly suggesting that either the septa had not yet been completed or that disassembly of the FtsZ polymers was compromised (Fig. 7). SflA and SflB reappeared at the interface between adjacent spores, perhaps to allow the disassembly of SepF, and hence destabilization of the Z-rings. The C-terminal part of SepF interacts with FtsZ in *B. subtilis* and *M. smegmatis* (Duman *et al*., 2013; Gola *et al*., 2015; Gupta *et al*., 2015; Hamoen *et al*., 2006). Though the Sfl proteins share significant homology with SepF in their C-terminal parts (Fig. 1), only SepF interacts with FtsZ (Schlimpert et al., 2017). Interestingly, in *Mycobacterium smegmatis* and *B. subtilis*, overexpression of SepF is lethal and largely blocks cell division, and it was suggested that this was due to interference of free SepF with the assembly of lateral cell division proteins (Gao *et al*., 2017; Gola *et al*., 2015). In *S. coelicolor* however, overexpression of SepF barely showed any effect, while overexpression of its paralogs SflA or SflB let to growth inhibited.

Taking into account the distinct localization patterns SepF and SflAB, and the *in vitro* interaction between these three proteins (Figure S4; (Schlimpert et al., 2017)), the activity of SflAB in terms of the disassembly of FtsZ filaments may be mediated via the disassembly of SepF rings. SflB lacks the N-terminal α-helix that is required for membrane lipid binding (Duman et al., 2013), suggesting that SflB will require SflA for membrane-specific localization. Extensive analysis using fluorescence microcopy showed that SflA and SflB foci are primarily formed before and after the cell division process, which is when SepF and FtsZ rings are formed. However, we occasionally observed colocalization of SflA or SflB with SepF (Fig. S5). Indeed, as discussed above, two-hybrid analysis also revealed interaction of SepF with the Sfl proteins. In the absence of SflAB, foci of FtsZ and SepF persist after spores have been formed, strongly suggesting that SflAB play a role in the termination of the cell division process.

We propose a model wherein SflA and SflB negatively affect the polymerization of SepF, thereby preventing the polymerization of SepF prior to the onset of cell division, and stimulating the depolymerization of SepF polymers after completion of cell division. During the onset of sporulation-specific cell division, SepF-rings assembly is initiated, initially whereby colocalizing with SflAB, which keep SepF inactive. Dispersal of SflAB then allows the formation of SepF rings, while SsgB localizes to recruit FtsZ, thus marking the start of cell division. Once completed, SflAB take up their positions again and assist in dispersing SepF, which leads to the destabilization of FtsZ filaments and their disassembly. The continued presence of SepF polymers in *sflAB* null mutants after the completion of sporulation-specific cell division would stabilize FtsZ filaments and continue to anchor them to the membrane, explaining why FtsZ polymers did not disassemble during spore maturation in these mutants.

Surprisingly, the localization of DivIVA was also disturbed in the aerial hyphae. While DivIVA is known to interact with a range of different protein partners, no interaction between DivIVA and either SepF or FtsZ has so far been reported (Halbedel and Lewis, 2019). Interestingly, DivIVA homologue GpsB was recently shown to interact with SepF in *Listeria monocytogenes* (Cleverley *et al*., 2019), while in *Staphylococcus aureus* GpsB was shown to interact with FtsZ to stimulate the formation of FtsZ bundles (Eswara *et al*., 2018). Biochemical experiments are required to establish how SflA and SflB affect the localization and/or polymerization of SepF, FtsZ and DivIVA.

Taken together, our work shows that SflAB control growth and cell division of the aerial hyphae of *Streptomyces*. Over-expression of the proteins strongly inhibits growth of the colonies, while in the absence of *sflA* and/or *sflB* DivIVA localizes ectopically, resulting in unusual branching of aerial hyphae. Besides controlling the localization and activity of DivIVA, SflAB also interact with - and control the localization of - SepF and hence of FtsZ. In the absence of SflAB, Z-rings and foci persist in mature spore chains. Thus SflAB ensure the correct localization of key cell division proteins in time and space during sporulation-specific cell division of *Streptomyces*.

## MATERIALS AND METHODS

### Bacterial strains and media

The bacterial strains used in this work are listed in Table S1. *E. coli* strains JM109 (Sambrook et al., 1989) and ET12567 (MacNeil et al., 1992) were used for routine cloning and for isolation of non-methylated DNA, respectively. *E. coli* transformants were selected on LB agar media containing the relevant antibiotics and grown O/N at 37°C. *Streptomyces coelicolor* A3(2) M145 was used as parental strain to construct mutants. All media and routine *Streptomyces* techniques are described in the *Streptomyces* manual (Kieser et al., 2000). Yeast extract-malt extract (YEME) and tryptic soy broth with 10% sucrose (TSBS) were the liquid media for standard cultivation. Regeneration agar with yeast extract (R2YE) was used for regeneration of protoplasts and with appropriate antibiotics for selection of recombinants (Kieser et al., 2000). Soy flour mannitol (SFM) agar plates were used to grow *Streptomyces* strains for preparing spore suspensions and for morphological characterization and microscopy.

### Plasmids and constructs and oligonucleotides

All plasmids and constructs described in this work are summarized in Table S2. The oligonucleotides are listed in Table S3.

#### Constructs for CRISPRi

As described previously, the 20 nt target sequence(spacer) was introduced into sgRNA scaffold by PCR using forward primers SepF_TF or SepF_NTF together with the reverse primer SgTermi_R_B (Ultee et al., 2020). The generated PCR products were cloned into pGWS1049 via restriction sites NcoI and BamHI to generate constructs pGWS1351 and pGWS1352. Subsequently, DNA fragments containing sgRNA scaffold and P*gapdh-dcas9* of constructs pGWS1049, pGWS1351 and pGWS1352 were digested with EcoRI and XbaI and cloned into pSET152 using the same restriction enzymes. The generated constructs pGWS1050 (no target sequence), pGWS1353(targeting template strand of *sepF*) and pGWS1354 (targeting non-template strand of *sepF*) were used in CRISPRi system.

#### Constructs for creating deletion mutants

Construction for in-frame deletion were based on the instable vector pWHM3 (Vara et al., 1989), essentially as described previously (Swiatek et al., 2012). For the deletion of *sflA*, its upstream region −1336/+9 (using primers sflA_LF-1339 and sflA _LR+9) and downstream region +427/+1702 (using primers sflA _RF_427 and sflA _RR+1702) were amplified by PCR from *S. coelicolor* M145 genomic DNA and cloned into pWHM3 as EcoRI-BamHI fragments, and the apramycin resistance cassette *aac(3)IV* flanked by *loxP* sites inserted in between. This resulted in plasmid pGWS750 that was used for deletion of *sflA* (SCO1749). The presence of *loxP* sites allows efficient removal of apramycin resistance cassette by Cre-recombinase (Fedoryshyn et al., 2008). The same strategy was used to create construct pGWS751 for the deletion of *sflB* (SCO5967). This plasmid contained the −1258/+9 and +357/+1917 regions relative to *sflB*, and the apramycin resistance cassette inserted in-between. The *sflA* and *sflB* double mutant (GAL16) was constructed in the background of a *sflA* in-frame deletion mutant (GAL14) by deleting *sflB*. For complementation of the *sflA* null mutant, pGWS1005 was used, an integrative vector based on pSET152 and harboring the entire coding region (+1/+468, amplified using primers sflA_F+1 and sflA_R+468) of *sflA* under control of the *ftsZ* promoter. Similarly, pGWS1006 was used for genetic complementation of *sflB* mutants, with pSET152 harboring the entire coding region (+1/+438, amplified using primers sflB-F+1 and sflB_R+438) of *sflB* under control of the *ftsZ* promoter.

#### Constructs for the expression of eGFP or E2-Crimson or dTomato fusion proteins

The eGFP gene was amplified by PCR from pKF41 using primers eGFP_F+1 and eGFP_R_717_Linker, adding a 12 bp linker in primer eGFP_R_717_Linker. The PCR fragments were digested with StuI and BamHI, and fused behind EcoRI and StuI digested fragment containing *ftsZ* promoter region excised from pGWS755 (Zhang et al., 2016). The fused EcoRI-StuI-BamHI fragment was then cloned into pSET152 via EcoRI and BamHI. Coding genes of *sflA* (amplified from *S. coelicolor* genomic DNA using primers sflA_F+1 and sflA_R+441), *sepF* (primers sepF_F+1 and sepF_R+639) and *sflB* (primers sflB_F+1 and sflB_R+411) were cloned in to the above construct via BamHI and XbaI to generate constructs pGWS784, pGWS785 and pGWS786, respectively. In pGWS786, the BamHI site between *egfp* and *sflB* was lost by fusion to the BglII site in PCR-amplified *sflB* DNA. The *E2-Crimson* gene was amplified by PCR from pTEC19 (Takaki et al., 2013) using primers E2Crimson_F_EEV and E2Crimson_linker_R_BH. The PCR fragment was digested with EcoRV and BamHI, and cloned into pGWS784 via StuI and BamHI to replace the gene for eGFP. Subsequently, the EcoRI -XbaI fragment containing P*_ftsZ_*-*E2-Crimson-sflA* was cloned into pHJL401 to generate pGWS1380. Similarly, *dTomato* gene was amplified by PCR from pLenti-V6.3 Ultra-Chili (Addgene plasmid # 106173) using primers dTomato_F_EEV and dTomato_linker_R_BH. The EcoRV and BamHI digested PCR was clone into StuI and BamHI digested pGWS785. Subsequently, the EcoRI-XbaI fragment containing *PftsZ-E2-dTomato-sepF* was cloned into pHJL401 to generate pGWS1383.

The coding region of *divIVA* (excluding the stop codon) together with its 393 bp upstream region were amplified by PCR from *S. venezuelae* genomic DNA using primers BglII-divIVA-SV-FW and NdeI-divIVA-SV-REV. The PCR product was cloned as BglII-NdeI fragment into pIJ8630 to generate construct pGWS800, which expresses DivIVA-eGFP under the control of the *divIVA* promoter.

#### Constructs for enhanced gene expression

To obtain enhanced expression of *sepF, sflA* and *sflB*, the genes were inserted behind the constitutive *ermE* promoter and an optimized ribosome binding site using plasmid pHM10a (Motamedi et al., 1995). For this, DNA fragments harboring the entire *sflA, sepF* or *sflB* coding region were amplified by PCR from *S. coelicolor* M145 genomic DNA using primer pairs *sflA*_F+4 and *sflA*_R+447, *sepF*_F+4 and *sepF*_R+648 and *sflB*_F+4 and *sflB*_R+417, respectively, and cloned into pHM10a digested with NdeI-HindII or NdeI-BamHI. The inserts of the pHM10a-based constructs were subsequently transferred as BglII-HindII or BglII-BamHI fragments to BamHI-HindII or BamHI digested pWHM3 to generate pGWS774 (for expression of *sflA*), pGWS775 (for *sepF*) and pGWS776 (for *sflB*).

#### Constructs for BACTH screening

The coding region of *sflA* was amplified from *S. coelicolor* M145 genomic DNA using primer pair *sflA-fw* and *sflA*-rv, and cloned as an XbaI-KpnI fragment into pUT18C and pKT25 to generate pBTH166 and pBTH167, respectively. *sepF* was amplified using primers *sepF-fw* and *sepF*-rv and cloned into pUT18C and pKT25 as an XbaI-XmaI fragment to generate pBTH110 and pBTH111, respectively. *sflB* was amplified from *S. coelicolor* M145 genomic DNA using primer pair *sflB-fw* and *sflB*-rv, cloned as an XbaI-KpnI fragment into pUT18C and pKT25, so as to generate pBTH170 and pBTH171, respectively. *sigR* was amplified from *S. coelicolor* M145 genomic DNA using primer pair SCO5216-fw and SCO5216-rv and cloned into pUC18 as an XbaI-XmaI fragment to generate pBTH17. Similarly, *rsrA* was amplified using primers SCO5217-fw and SCO5217-rv and cloned into pKT25 as XbaI-XmaI fragment to generate pBTH23.

### Microscopy

#### Light microscopy

Sterile cover slips were inserted at an angle of 45 degrees into SFM agar plates, and spores of *Streptomyces* strains were carefully inoculated at the intersection angle. After incubation at 30°C for 3 to 5 days, cover slips were positioned on a microscope slide prewetted 5 μl of 1xPBS. Fluorescence and corresponding light micrographs were obtained with a Zeiss Axioscope A1 upright fluorescence microscope (with an Axiocam Mrc5 camera at a resolution of 37.5 nm/pixel). The green fluorescent images were created using 470/40 nm band pass (bp) excitation and 525/50 bp detection, for the red channel 550/25 nm bp excitation and 625/70 nm bp detection was used (Willemse and van Wezel, 2009). DAPI was detected using 370/40 nm excitation with 445/50 nm emission band filter. For staining of the cell wall (peptidoglycan) we used FITC-WGA, for membrane staining FM5-95 and for DNA staining DAPI (all obtained from Molecular Probes). For stereomicroscopy we used a Zeiss Lumar V12 stereomicroscope. All images were background corrected setting the signal outside the hyphae to zero to obtain a sufficiently dark background. These corrections were made using Adobe Photoshop CS4.

#### Electron microscopy

Morphological studies on surface grown aerial hyphae and/or spores by cryo-scanning electron microscopy were performed using a JEOL JSM6700F scanning electron microscope as described previously (Colson et al., 2008). Transmission electron microscopy (TEM) for the analysis of cross-sections of hyphae and spores was performed with a FEI Tecnai 12 BioTwin transmission electron microscope as described (Piette et al., 2005).

### BATCH complementation assay

For BACTH complementation assays, vectors pKT25 and pUT18C harboring genes of interest were used in various combinations to co-transform *E. coli* BTH101 cells carrying plasmid pRARE (Novagen). The transformants were plated onto LB medium containing ampicillin (100 μg/mL), kanamycin (50 μg/mL) chloramphenicol (50 μg/mL) and were incubated for 24-36 h at 30°C. Then 3 independent representative co-transformants were grown on M63 minimal medium agar plates containing proper antibiotics ampicillin 50 μg/ mL, kanamycin 25 μg/ mL and chloramphenicol 25 μg/ mL. This medium allows growth of co-transformants only if the co-expressed proteins interact with each other. Co-transformation of pBTH17 (*sigR*) and pBTH23 (*rsrA*) was used as positive control, while co-transformation of empty plasmids pUT18 and pKT25 was used as negative control.

### Computer analysis

For DNA and protein searches used StrepDB (http://strepdb.streptomyces.org.uk/) and STRING (http://string.embl.de). Alignment was built using Clustal Omega (http://www.ebi.ac.uk/Tools/msa/clustalo/) and Boxshade program. Secondary structures of proteins were predicted using JPRED (http://www.compbio.dundee.ac.uk/jpred4/index_up.html).

## Supporting information

All Supplemental Information

