## Supplementary material for "Branching of sporogenic aerial hyphae in *sflA* and *sflB* mutants of *Streptomyces coelicolor* correlates to ectopic localization of DivIVA and FtsZ in time and space": All Supplemental Information

Belonging to the manuscript:

**Table S1.** Bacterial strains

| Bacteria strains | Genotype | Reference |
|---|---|---|
| *E. coli* JM109 | See reference | (Sambrook et al., 1989) |
| *E.coli* ET12567 | See reference | (MacNeil et al., 1992) |
| *S. coelicolor* M145 | SCP1$^-$ SCP2$^-$ | (Kieser et al., 2000) |
| K202 | M145 + KF41 | (Grantcharova et al., 2005) |
| GAL14 | M145Δ*sflA* | This work |
| GAL15 | M145Δ*sflB* | This work |
| GAL16 | M145Δ*sflA*Δ*sflB* | This work |
| GAL18 | M145 + pGWS760 | This work |
| GAL111 | M145 + pGWS784 | This work |
| GAL112 | M145 + pGWS785 | This work |
| GAL113 | M145 + pGWS786 | This work |
| GAL44 | M145 + pGWS774 | This work |
| GAL45 | M145 + pGWS775 | This work |
| GAL46 | M145 + pGWS776 | This work |
| GAL70 | M145 + pWHM3 | This work |
| GAL123 | GAL14 + pKF41 | This work |
| GAL124 | GAL15 + pKF41 | This work |
| GAL125 | M145 + pGWS800 | This work |
| GAL126 | GAL14 + pGWS800 | This work |
| GAL127 | GAL15 + pGWS800 | This work |
| GAL128 | GAL14 + pGWS1005 | This work |
| GAL129 | GAL15 + pGWS1006 | This work |
| GAL130 | M145 + pGWS1351 | This work |
| GAL131 | M145 + pGWS1352 | This work |
| GAL132 | M145 + pGWS1353 | This work |
| GAL133 | M145 + pGWS785 + pGWS1380 | This work |
| GAL134 | M145 + pGWS786 + pGWS1383 | This work |
| GAL135 | M145 + pGWS786 + pGWS1380 | This work |

**Table S2.** Plasmids and Constructs

| Plasmid and constructs | Description | Reference |
|---|---|---|
| pWHM3 | *E. coli*/*Streptomyces* shuttle vector, multi-copy and very unstable in *Streptomyces* | (Vara et al., 1989) |
| pSET152 | *E. coli*/*Streptomyces* shuttle vector, high copy number in *E. coli* and integrative in *Streptomyces* | (Bierman et al., 1992) |
| pHJL401 | E. coli/Streptomyces shuttle vector, around five copies per chromosome in Streptomyces and around 100 copies per chromosome in E. coli. | (Larson and Hershberger, 1986) |
| pKF41 | Integrative construct expressing ftsZ-egfp from the natural ftsZ promoter region | (Grantcharova et al., 2005) |
| pHM10a | Conjugative E. coli-Streptomyces shuttle vector, harboring PermE and an optimized RBS | (Motamedi et al., 1995) |
| pIJ8630 | *E. coli*/*Streptomyces* shuttle vector, containing the promoterless eGFP and integrative in *Streptomyces* | (Sun et al., 1999) |
| pGWS750 | pWHM3 containing flanking regions of *S. coelicolor* SCO1749 with apraloxP-*Xba*I inserted between then in pWHM3 EcoRI-HindIII | This work |
| pGWS751 | pWHM3 containing flanking regions of *S. coelicolor* SCO5967 with apraloxP-*Xba*I inserted between then in pWHM3 EcoRI-HindIII | This work |
| pUWL-Cre | Cre-recombinase expression plasmid | (Fedoryshyn et al., 2008) |
| pUT18 | BATCH plasmid containing the T18 fragment of *cya* gene, fused behind multiple cloning site. | (Karimova et al., 1998) |
| pUT18C | BATCH plasmid containing the T18 fragment of *cya* gene, followed by multiple cloning site. | (Karimova et al., 1998) |
| pKT25 | BATCH plasmid containing the T25 fragment of *cya* gene, followed by multiple cloning site. | (Karimova et al., 1998) |
| pBTH17 | pUT18 containing +1/+681 region of *sigR* (SCO5216) from *S .coelicolor* fused in front of the *cya* gene. | This work |
| pBTH23 | pKT25 containing +1/+315 region of *rsrA* (SCO5217) from *S. coelicolor* fused behind the cya gene. | This work |
| pBTH166 | pUT18C harboring +1/+438 region of *sflA* from *S. coelicolor* fused behind the *cya* gene | This work |
| pBTH110 | pUT18C harboring +1/+639 region of *sepF* from *S. coelicolor* fused behind the *cya* gene | This work |
| pBTH170 | pUT18C harboring +1/+408 region of *sflB* from *S. coelicolor* fused behind the *cya* gene | This work |

| | | |
|---|---|---|
| pBTH167 | pKT25 harboring +1/+438 region of *sflA* from *S. coelicolor* fused behind the *cya* gene | This work |
| pBTH111 | pKT25 harboring +1/+639 region of *sepF* from *S. coelicolor* fused behind the *cya* gene | This work |
| pBTH171 | pKT25 harboring +1/+408 region of *sflB* from *S. coelicolor* fused behind the *cya* gene | This work |
| pGWS784 | pSET152 harboring *egfp-sflA* under control of *ftsZ* promoter | This work |
| pGWS785 | pSET152 harboring *egfp-sepF* under control of *ftsZ* promoter | This work |
| pGWS786 | pSET152 harboring *egfp-sflB* under control of *ftsZ* promoter | This work |
| pGWS774 | pWHM3 containing P*ermE*, RBS and *sflA* | This work |
| pGWS775 | pWHM3 containing P*ermE,* RBS and *sepF* | This work |
| pGWS776 | pWHM3 containing P*ermE,* RBS and *sflB* | This work |
| pGWS800 | pIJ8630 expressing *S. venezuelae* DivIVA(SVEN_1732)-eGFP from its native promoter | This work |
| pGWS1005 | pSET152 harboring the +1/+468 region of *sflA* under the control of *ftsZ* promoter of *S. coelicolor* | This work |
| pGWS1006 | pSET152 harboring the +1/+438 region of *sflB* under the control of *ftsZ* promoter of *S. coelicolor* | This work |
| pGWS1049 | pHJL401 containing sgRNA scaffold (no scpacer) and *dcas9* under the control of *gapdh* promoter | (Ultee et al., 2020) |
| pGWS1050 | pSET152 containing sgRNA scaffold (no scpacer) and *dcas9* under the control of *gapdh* promoter | This work |
| pGWS1351 | pHJL401 containing sgRNA scaffold with spacer targeting template strand of *sepF* and *dcas9* under the control of *gapdh* promoter | This work |
| pGWS1352 | pHJL401 containing sgRNA scaffold with spacer targeting non-template strand of *sepF* and *dcas9* under the control of *gapdh* promoter | This work |
| pGWS1353 | pSET152 containing sgRNA scaffold with spacer targeting template strand of *sepF* and *dcas9* under the control of *gapdh* promoter | This work |
| pGWS1354 | pSET152 containing sgRNA scaffold with spacer targeting non-template strand of *sepF* and *dcas9* under the control of *gapdh* promoter | This work |
| pGWS1380 | pHJL401 harboring E2-Crimson-sflA under control of ftsZ promoter | This work |
| pGWS1383 | pHJL401 harboring dTomato-sepF under control of ftsZ promoter | This work |

**Table S3.** Oligonucleotides

| Name | 5'-3' sequence[#] |
|---|---|
| sflA_LF -1336 | GTCA**GAATTC**GTTGAAGGTGCCGCAGCACATCTG |
| sflA_LR +9 | GTCAGAAGTTATCCATCACC**TCTAGA**CGATCCCATGGACGCCTCCTCTCA |
| sflA_RF+427 | GTCAGAAGTTATCGCGCATC**TCTAGA**TTCAACCAGAGCTGAGGCGGGGCC |
| sflA_RR +1702 | GTCA**AAGCTT**CCCATGGCCGCGTCCCCGAAGTT |
| sflB_LF-1258 | GTCA**GAATTC**GAACTGCACCATCAGGTAGGCGT |
| sflB_LR+9 | GTCAGAAGTTATCCATCACC**TCTAGA**CGATTTCACTCGCCTTCATTGCCTGCA |
| sflB_RF +357 | GTCAGAAGTTATCGCGCATC**TCTAGA**ACGTCTTCCTGCTGACCCCGGCG |
| sflB_RR+1917 | GTCA**AAGCTT**CGGTCACGGGCTCAGGTGTACCA |
| sflA_F +1 | GTCAGAATTC**AGGCCT**TCGACATGGGATCGGTACGCAAGGCG |
| sflA_R+468 | gcta**GGATCC**TGCGGGGCCGTGGTGTGCGGC |
| sflB_F +1 | GTCAGAATTC**AGGCCT**TCGACGTGAAATCGGGGGAGCCGGTG |
| sflB_R+438 | gtca**AGATCT**CCGGACGGTCAGGGCCGGAGC |
| BglII-divIVA-SV-FW | GAC**AGATCT**GGCCGGACAAGCGAGAGCACG |
| NdeI-divIVA-SV-REV | GAC**CATATG**GTTGTCGTCCTCGTCGATCAGG |
| sepF_F +1 | GTCAGAATTC**AGGCCT**TCGACATGGCCGGCGCGATGCGCAAG |
| sepF_R +639 | GTCAAAGCTT**GGATCC**CTCTGGTTGAAGAACCCGCC |
| eGFP_F+1_ES | CTGAGAATTC**AGGCCT**TCGACATGGTGAGCAAGGGCGAGGAGCTGT |
| eGFP_R+717_linker_BH | CTGAAAGCTT**GGATCC**GGTGGCGACCGGCTTGTACAGCTCGTCCATGCCGAGA |
| E2Crimson_F_EEV | CTGAGAATTC**GATATC**TCGACATGGATAGCACTGAGAACGTCATC |
| E2Crimson_linker_R_BH | CTGAAAGCTT**GGATCC**GGTGGCGACCGGCTGGAACAGGTGGTGGCG |
| dTomato_F_EEV | CTGAGAATTC**GATATC**TCGACATGGTGAGCAAGGGCGAG |
| dTomato_linker_R_BH | CTGAAAGCTT**GGATCC**GGTGGCGACCGGCTTGTACAGCTCGTCCATGCC |
| sflA_F+1_EB | CTGAGAATTC**GGATCC**ATGGGATCGGTACGCAAGGCGA |
| sflA_R+441_XH | CTGAAAGCT**TCTAGA**TCAGCTCTGGTTGAAGAATC |
| sflB_F+1_EBg | CTGAGAATTC**AGATCT**GTGAAATCGGGGGAGCCGGTGA |
| sflB_R+411_XH | CTGAAAGC**TCTAGA**TTACACTCCCGGCACCCCCGC |
| sflA_F+4 | GCTAGAATT**CATATG**GGATCGGTACGCAAGGCGAGT |
| sflA_R+447 | GCTA**AAGCTT**CCCGCCTCAGCTCTGGTTGAA |
| sepF_F+4 | GTCAGAATT**CATATG**GCCGGCGCGATGCGCAAGATG |
| sepF_R+648 | GTCAAAGCTT**GGATCC**TAGTGCGTCTCAGCTCTGGTT |
| sflB_F+4 | GTCAGAATT**CATATG**AAATCGGGGGAGCCGGTGAAC |
| sflB_R+417 | GTCA**AAGCTT**GGACCGTCACACTCCCGGCAC |
| sflA-fw | CGC**TCTAGA**CATGGGATCGGTACGCAAGGC |
| sflA-rv | CGG**GGTACC**CAGCTCTGGTTGAAGAATCCG |
| sepF-fw | CG**TCTAGA**CATGGCCGGCGCGATGCGC |
| sepF-rv | CT**CCCGGG**AGCTCTGGTTGAAGAACCCGCC |
| sflB-fw | CGC**TCTAGA**AGTGAAATCGGGGGAGCCGGT |
| sflB-rv | CGG**GGTACC**CACACTCCCGGCACCCCCGCC |
| SCO5216-fw | CGC**TCTAGA**GATGGGTCCGGTCACTGGG |
| SCO5216-rev | CGC**CCCGGG**ATGACCCCGAGCCTTTCGC |
| SCO5217-fw | CCT**TCTAGA**CATGAGCTGCGGAGAGCCG |
| SCO5217-rev | GCA**CCCGGG**AGGACTCCTGCGGGGCCGAC |
| SgTermi_R_B | ctag**GGATCC**CAAAAAACCCCTCAAGACCCGTTTAGAGGCCCCAAGGGGTTATGC TAGTTACGCCTACGTAAAAAAAGCACCGACTCGGTGCC |
| SepF_TF | CATG**CCATGG**TGTCCGAACGAGAGCCCTACGTTTTAGAGCTAGAAATAGC |
| SepF_NTF | CATG**CCATGG**CGTACCCATCGTCCTCCACGGTTTTAGAGCTAGAAATAGC |

[#] Restriction sites used for cloning are underlined and in bold. GGATCC, BamHI; AGATCT, BglII; GAATTC, EcoRI; GATATC, EcoRV; AAGCTT, HindIII; GGTACC, KpnI; CATATG, NdeI; CCATGG, NcoI; AGGCCT, StuI; TCTAGA, XbaI; CCCGGG, XmaI. T7 terminator in primer SgTermi_R_B and *aph* terminator in primer Cas9Termi_R+4107_XH are shown in red. The 20 nt target in primers SepF-TF and SepF_NTF ares shown in blue.

**SUPPLEMENTAL FIGURES**

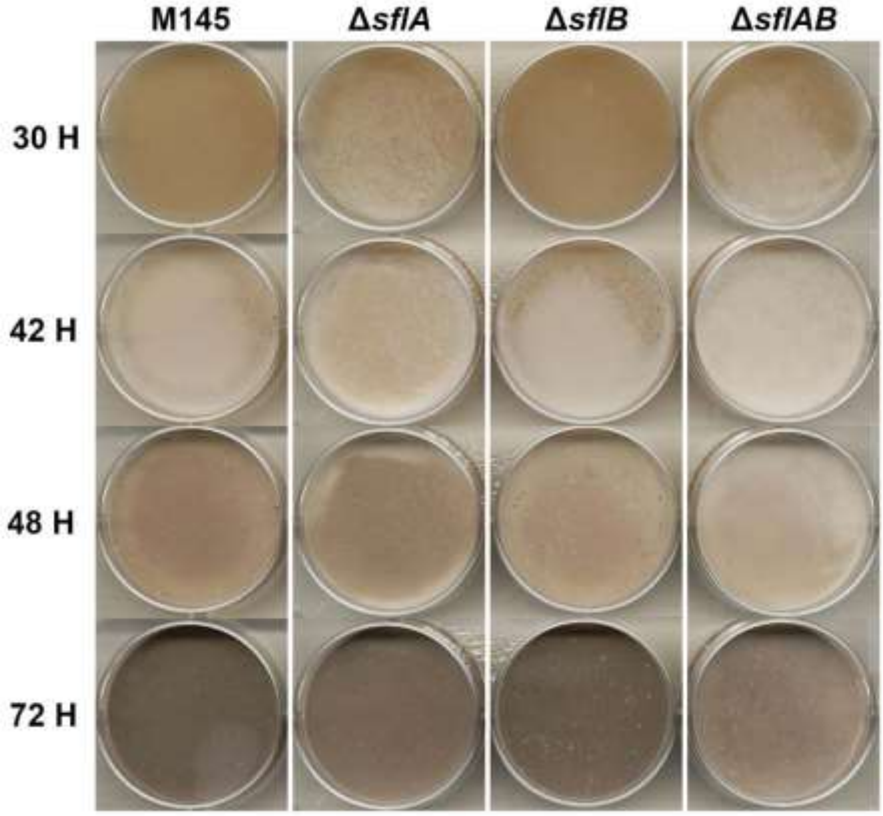


**Figure S1. Growth of *S. coelicolor* M145 and its *sfl* mutants on SFM medium at different time points.** Spores of *S. coelicolor* and its *sfl* mutants were counted using haemocytometer, and the titer in terms of colony forming units (cfu/mL) were determined. The same amount of viable spores (10E+05) were plated onto SFM agar on 6-well plate at 30°C and the plates were scanned in the incubator at 30 min intervals. *sflA* and *sflAB* mutants produced abundant white-pigmented aerial hyphae after 30 hours incubation, when the parent and *sflB* mutant were still in vegetative growth. At 42 hours, *sflA* mutant already entered sporulation, as indicated by the grey pigmentation, while the parent and other mutants had now initiated aerial growth. At 48 hours, *sflB* and *sflAB* mutants just started sporulation while wild type and *sflA* mutant had produced decent amount of grey spores . After 72 hours, all strains had fully developed though *sflA* and *sflAB* mutants showed slightly lighter grey color suggesting probably less spores formation.

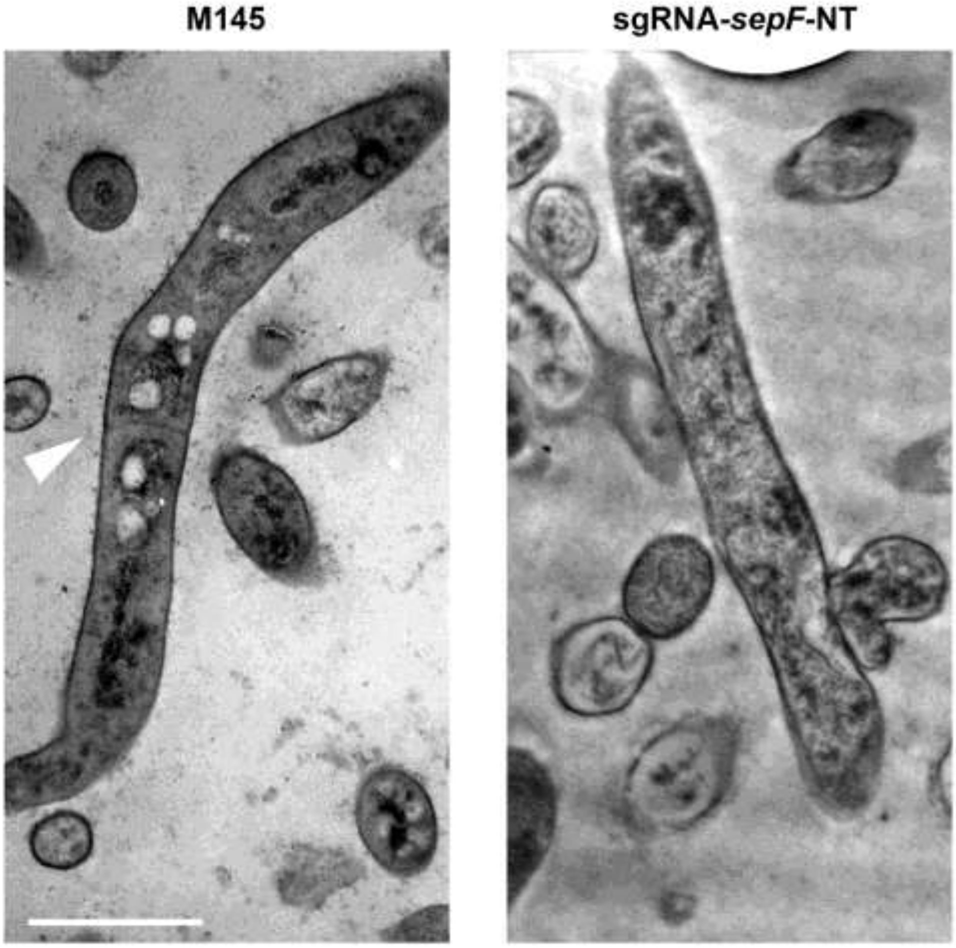


**Figure S2. Transmission electron micrographs of *S. coelicolor* M145 and *sepF* knockdown mutant.** Cross wall in vegetative hyphae wild type M145 was synthesized normally, resulted into compartments containing multiple chromosomes. When non-template strand of *sepF* was targeted by CRISPRi system(sgRNA-sepF-NT), cross-wall formation was nearly abolished and chromosomal DNA became more dispersed. Cultures were grown on SFM agar plates for 5 days at 30$^{0}$C. Arrow indicates cross-wall in wild type M145. Bar, 1 µm.

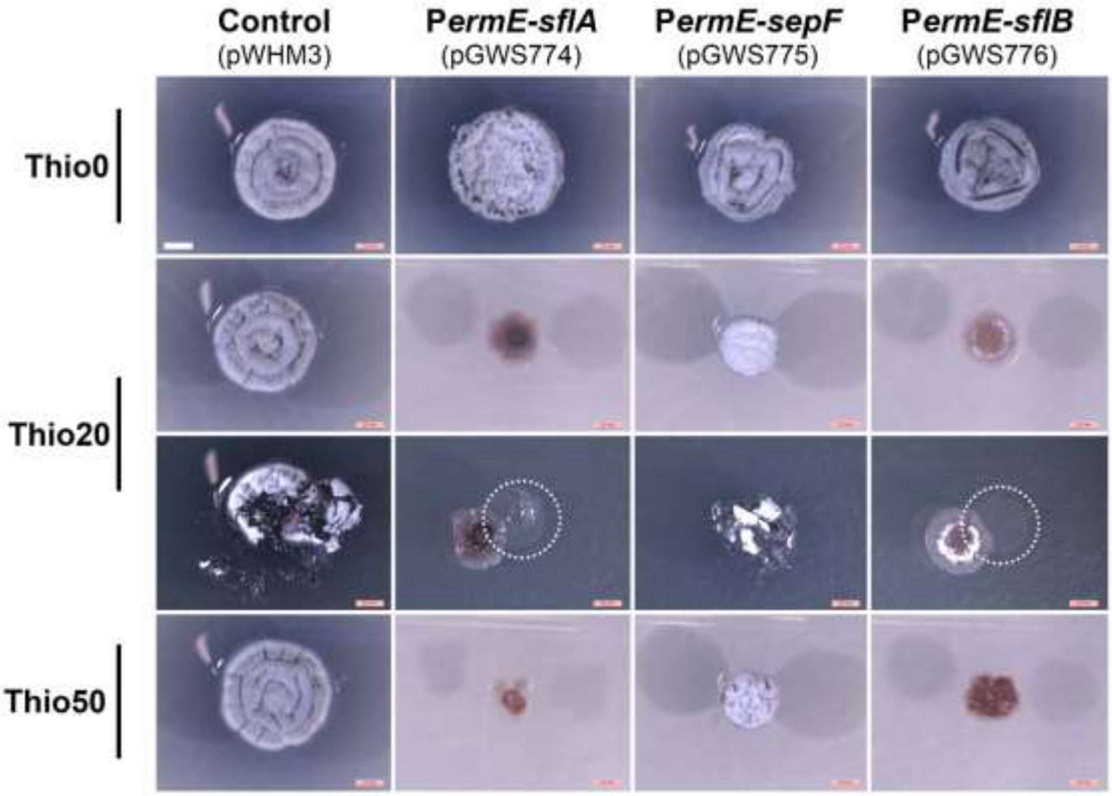


**Figure S3. Effect of enhanced expression of *sepF* and *sfl* genes on colony morphology.** On R5 agar plates without thiostrepton, all colonies developed normally and had comparable colony sizes. However, in the presence of thiostrepton (20 µg/ml), the colonies harboring the SflA and SflB expression constructs were blocked in development, while those expressing SepF produced aerial hyphae but formed significantly smaller colonies, indicative of growth inhibition. Interestingly, colonies overexpressing SflA (GAL44) or SflB (GAL45) did not attach to the agar surface, whereby colonies could be readily mashed up or moved over the agar surface with a toothpick. In contrast, wild-type colonies or colonies expressing SepF (GAL45) were solid and firmly attached to the agar surface. When the thiostrepton concentration was increased further to 50 µg/mL, colonies of GAL44 and GAL46 more or less floated on the agar surface, while those of GAL70 and GAL45 were still firmly attached to the agar surface (not shown). In particular GAL44 produced minute colonies, indicative of strong retardation of growth by the enhanced expression of SflA.

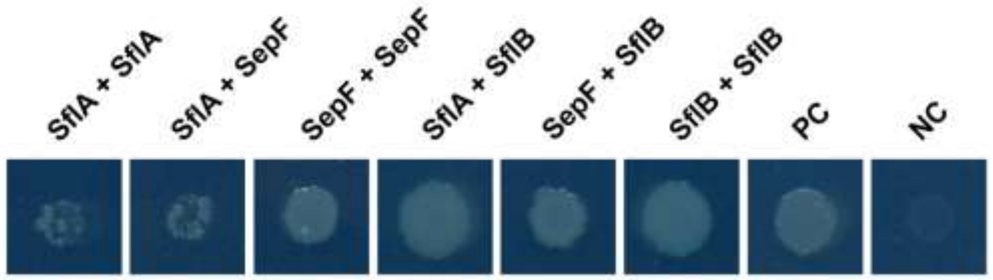


**Figure S4. Analysis of protein interaction using the bacterial two-hybrid system.** Possible interactions between SflA, SflB and SepF from *S. coelicolor* were assayed on M63 agar plates containing proper antibiotics for 4 days. Growth indicates interaction between the partners. All three SepF-like proteins interacted with each other and all showed self-interaction.

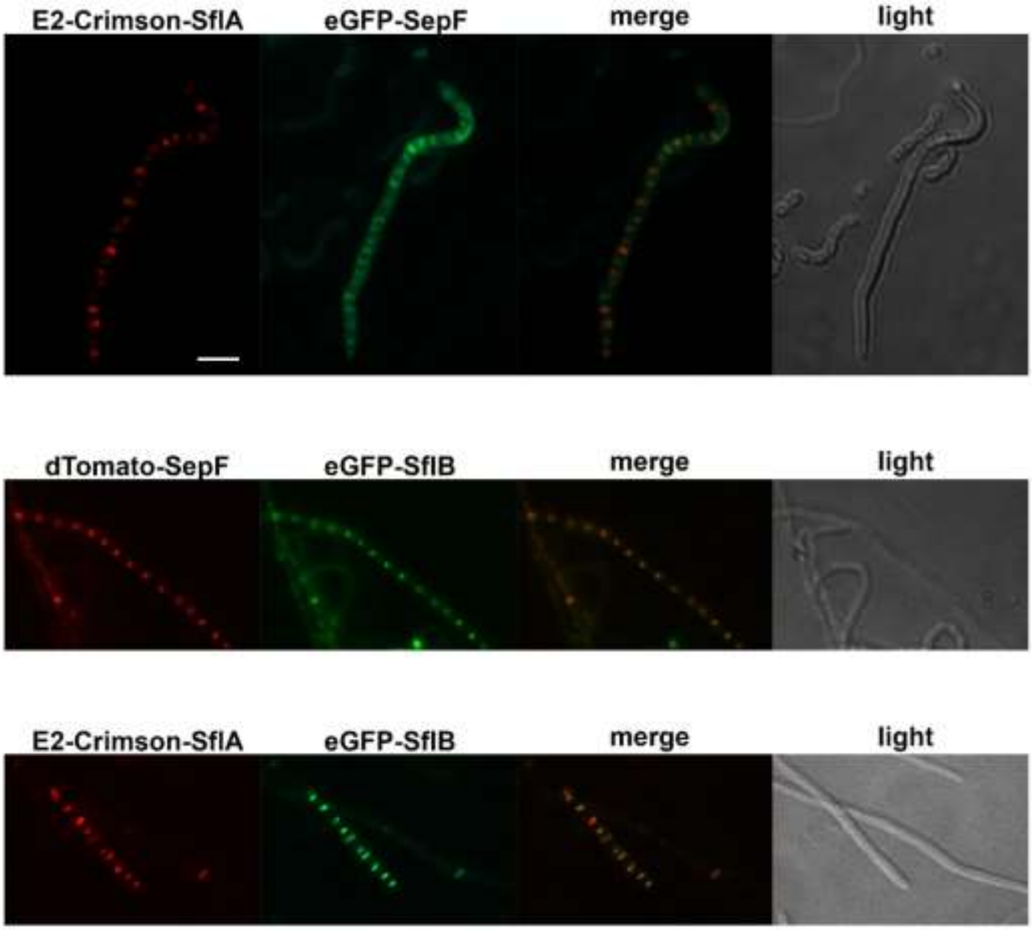


**Figure S5. Colocalization studies of SflA, SflB and SepF**. Sporogenic aerial hyphae were imaged by fluorescence microscopy visualizing the eGFP fusion proteins (green), E2-Crimson or dTomato fusion proteins (red). Corresponding light micrographs are shown on the right. Bar, 2 µm.
